# A Cancer-Specific Antigen Drives Histone Acetylation by Stabilizing the Acetyltransferases

**DOI:** 10.1101/2025.03.05.641767

**Authors:** Xuekun Fu, Xu Yang, Jie Huang, Patrick Ryan Potts, Chao Liang

**Affiliations:** Department of Systems Biology, School of Life Sciences, Southern University of Science and Technology, Shenzhen, 518055, China; Department of Computational Biology, St. Jude Children’s Research Hospital, Memphis, TN, 38105, USA; Institute of Integrated Bioinfomedicine and Translational Science (IBTS), School of Chinese Medicine, Hong Kong Baptist University, Hong Kong SAR, 999077, China; Department of Cell and Molecular Biology, St. Jude Children’s Research Hospital, Memphis, TN, 38105, USA

## Abstract

Histone acetylation is a critical epigenetic modification that regulates gene expression by modulating chromatin structure and function. Histone acetyltransferases are essential for maintaining acetylation homeostasis and the disruption of this balance can lead to aberrant gene expression and cancer development. Here, we describe that the cancer-specific protein MAGE-A10 increases cellular histone acetylation by stabilizing essential histone acetyltransferases KAT2A and KAT2B. The aberrant expression of MAGE-A10 in tumors prevents the degradation of KAT2A/2B through p62-mediated autophagy. Mechanistically, MAGE-A10 antagonizes the binding of KAT2A/2B with the E3 ubiquitin ligase complex CUL4A-DDB1, thereby decreasing the formation of their K63-linked ubiquitination. Furthermore, KAT2A enhances the transcription of the MAGE-A10 gene, forming a positive feedback loop that contributes to tumorigenesis. These findings provide insights into the molecular mechanisms hijacked in cancer that drive perturbed histone acetylation and suggest potential therapeutic strategies.

## INTRODUCTION

Histone acetylation is a critical epigenetic modification that regulates gene expression by altering chromatin structure and function^1–3^. This process leads to chromatin changes that can influence gene expression and affect numerous biological functions, including the development of various diseases such as cancers^3–6^. The dynamic balance of histone acetylation is maintained by the opposing activities of histone acetyltransferases (HATs) and histone deacetylases (HDACs)^7,8^. Among the HATs, KAT2A (also known as GCN5) and KAT2B (also known as PCAF) play particularly significant roles. These enzymes primarily acetylate lysine residues on histone tails, specifically H3K9 and H3K14, resulting in transcriptional activation that is crucial for various cellular processes, including DNA repair, cell cycle progression, and transcriptional regulation^9–13^.

The proper function and regulation of KAT2A and KAT2B are crucial for maintaining acetylation homeostasis, and their dysregulation has been implicated in the development and progression of cancer^14–16^. Notably, the high incidence of mutations and the pivotal roles of KAT2A/2B in cancer development have garnered significant attention^17–20^. Elevated KAT2A/2B expression is associated with poor prognosis in breast cancer, non-small cell lung carcinoma, melanoma, renal cell carcinoma, acute myeloid leukemia and colon cancer^20–23^. This is primarily due to histone acetylation-mediated co-activation of E2F and MYC transcriptional targets, which sustain cell proliferation and survival^24,25^. Understanding the mechanisms that control the stability and activity of these acetyltransferases is, therefore, vital for elucidating their roles in tumorigenesis and for identifying potential therapeutic targets.

Melanoma antigen (MAGE) genes encode a family of proteins characterized by a common MAGE homology domain (MHD)^26–28^. Most human MAGE proteins, including MAGE-A10, are classified as cancer-testis antigens (CTAs) due to their physiological restriction to the testis and aberrant expression in various cancers^28–33^. While the precise mechanisms regulating MAGE expression are not fully understood, MAGE CTAs have garnered increasing interest as hallmarks of cancer, owing to their widespread expression in aggressive tumors, association with poor clinical prognosis, and their oncogenic potential to enhance tumor growth and metastasis^34–36^. A notable mechanism of MAGE proteins involves their role as adaptors for E3 ubiquitin ligases, targeting novel proteins for degradation^37–39^. Despite this, the precise molecular mechanisms and oncogenic potential of many MAGE CTAs, including MAGE-A10, remain largely unknown.

Here, we demonstrate that the normally germ-cell-restricted MAGE-A10 is aberrantly expressed in cancer and functions as a potent oncogene, driving tumorigenesis by stabilizing KAT2A and KAT2B expression. MAGE-A10 acts as a binding competitor for the CUL4A-DDB1 E3 ubiquitin ligase complex, preventing K63-linked ubiquitination of KAT2A/2B in cancer cells. This antagonistic effect suppresses p62-mediated autophagic degradation of KAT2A/2B, leading to increased cellular histone acetylation and downstream oncogene expression. Furthermore, stabilized KAT2A enhances the transcription of MAGE-A10, forming a positive feedback loop. These findings highlight the critical role of MAGE-A10 in disrupting normal cellular regulatory mechanisms to promote cancer progression.

## RESULTS

### MAGE-A10 Is Aberrantly Expressed in Cancer and Is Necessary and Sufficient to Drive Tumor Growth

To comprehensively examine the expression pattern of *MAGEA10*, we performed RNA-seq analysis on 29 non-diseased human tissues from the GTEx project and observed that their expression is predominantly restricted to the testis (**Figure 1A**). Consistent with other CTA genes, *MAGEA10* is aberrantly expressed in various tumors. Our analysis of transcriptomic data from The Cancer Genome Atlas (TCGA) revealed that *MAGEA10* is frequently expressed in multiple cancer types, including bladder urothelial carcinoma (BLCA), skin cutaneous melanoma (SKCM), and lung squamous cell carcinoma (LUSC) (**Figure 1B**). Additionally, immunohistochemistry staining of SKCM and LUSC specimens confirmed the presence of MAGE-A10 protein in tumor tissues (**Figure S1A**).

**Figure 1.**
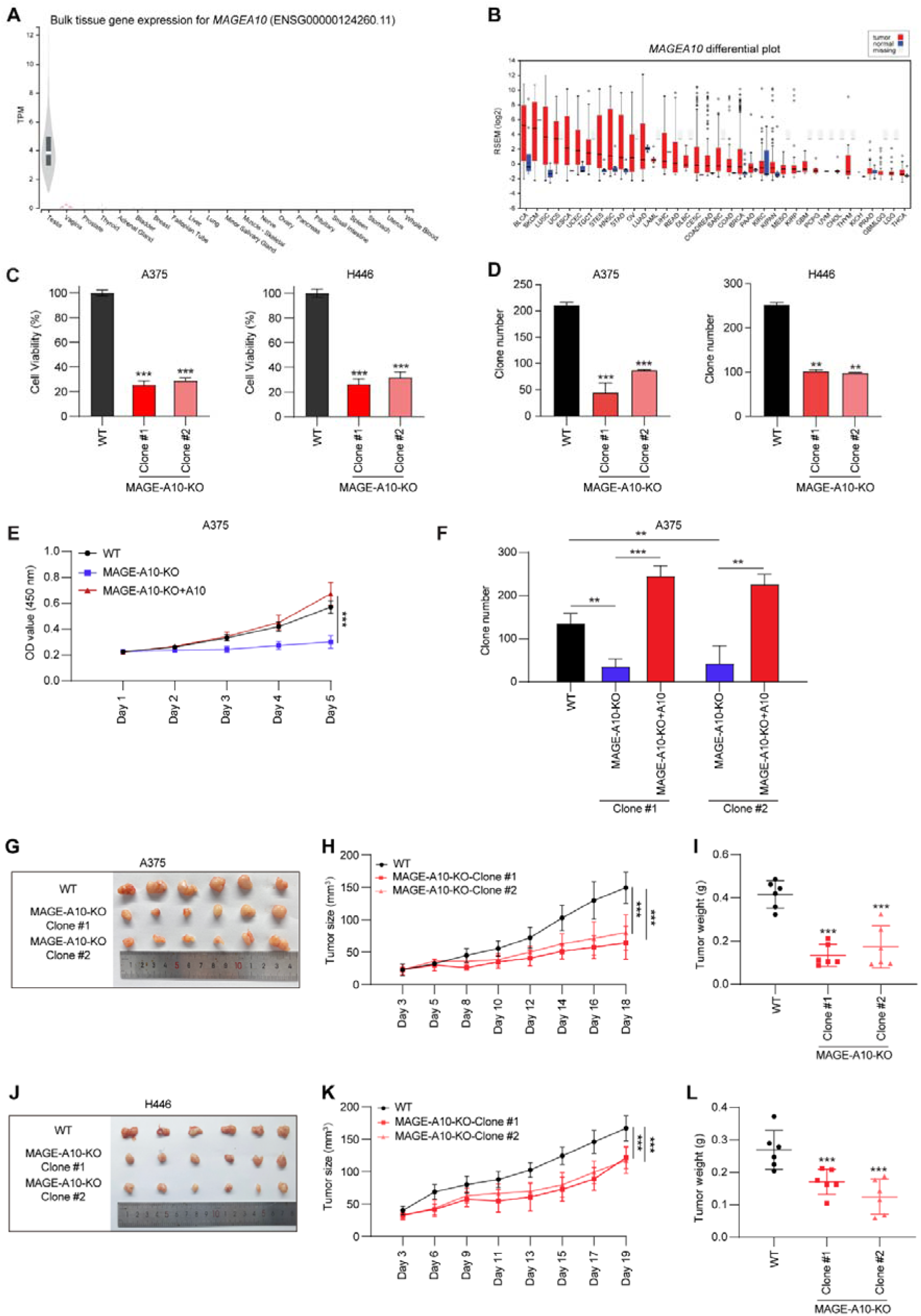
MAGE-A10 Is Aberrantly Expressed in Cancer and Is Necessary and Sufficient to Drive Tumor Growth. (A) RNA-Seq data from GTEx of the normalized expression of human *MAGEA10* in the indicated tissues. The numbers of biological replicates are as follows: testis 361, brain 255, small intestine 187, white adipose 663, muscle 803, skin 604, esophagus 555, stomach 359, liver 226, heart 429. blood vessel 663, lung 578, colon 373, nerve 619, pituitary 283, blood 755, adrenal gland 258, kidney 85, prostate 245, salivary gland 162, ovary 180, breast 459, pancreas 328, vagina 156, uterus 142, spleen 241, fallopian tube 9, bladder 21, cervix 10. Central band indicates median, boxes define 25 and 75 percentiles, and whiskers define 5 and 95 percentiles. (B) Expression of *MAGEA10* in normal adjacent (blue) or tumor tissue (red) from TCGA RNA-Seq data. N.D. represents normal adjacent is not available. Data visualized by firebrowse. Number of biological replicates are as follows: ACC 41, BLCA 307, BRCA 524, CESC 175, CHOL 11, COAD 153, COADREAD 232, DLBC 36, ESCA 121, HNSC 416, KICH 46, KIPAN 515, KIRC 411, KIRP 58, LAML 7, LGG 528, LUAD 301, LUSC 400, MESO 31, OV 183, PCPG 70, PRAD 204, READ 79, SARC 156, SKCM 422, STAD 288, STES 409, TGCT 126, THCA 408, THYM 75, UCEC 306, and UCS 37. Central band indicates median, boxes define 25 and 75 percentiles, and whiskers define 5 and 95 percentiles. (C) The CCK-8 assay was performed to evaluate the viability of Wild-type or MAGE-A10-knockout A375 and H446 cells. (D) Wild-type or MAGE-A10-knockout A375 and H446 clones were assayed for clonogenic growth. (E) MAGE-A10-knockout A375 cells or those reconstituted with MAGE-A10 were counted for cell proliferation by CCK8 assay at the indicated time points (n = 6 biological replicates). (F) Re-expression of MAGE-A10 rescues clonogenic growth of MAGE-A10-knockout A375 cells. (G - L) Knockout of MAGE-A10 in A375 (G-I) or H446 (J-L) decreases xenograft tumor growth in mice (n = 6 per. group). Data are means ± SDs. Asterisks indicate significant differences from the control (*p<0.05, **p<0.01, ***p<0.001, n.s., not significant).

To ascertain whether the aberrant expression of MAGE-A10 in tumor cells is merely a passenger event resulting from global genomic dysregulation or if MAGE-A10 actively contributes to tumorigenesis, we conducted a series of gain- and loss-of-function studies to elucidate the role of MAGE-A10 in promoting cancer cell growth. First, we examined whether multiple cancer cells require the expression of MAGE-A10 for viability. Intriguingly, knockout of MAGE-A10 in A375, SK-MEL-2 melanoma cells, H446 small cell lung cancer cells, and M059K glioblastoma cells resulted in dramatic decrease in cell viability (**Figure 1C**) and clonogenic growth (**Figure 1D** and **S1B**). Furthermore, knockout of MAGE-A10 decreased the proliferation rate and clonogenic growth of A375 and H446 cells, which could be rescued by re-expression of MAGE-A10 (**Figure 1E-F** and **S1C–S1E**). Consistent with these findings, knockout of MAGE-A10 slowed xenograft tumor growth (**Figures 1G-L** and **S1F**). Finally, to determine whether overexpression of MAGE-A10 is sufficient to drive tumorigenic phenotypes, we stably expressed MAGE-A10 in H460, H1650, H1975 non-small cell lung cancer cells, Hela cervical cancer cells that do not naturally express MAGE-A10 or M8 melanoma cells. Strikingly, expression of MAGEA10 accelerated the growth in these cells (**Figure S1G**) and xenograft tumor growth of H1975 cells and M8 melanoma cells in mice (**Figures S1H and S1I**). Together, these results suggest that MAGE-A10 is normally restricted to expression in the testis but is aberrantly expressed in a variety of cancers, where it is necessary and sufficient to drive tumorigenesis.

### MAGE-A10 Promotes KAT2A/KAT2B Expression and Histone 3 Lysine Acetylation

To elucidate the molecular mechanisms of MAGE-A10 oncogenic activity, we performed unbiased analysis of MAGE-A10 interacting proteins by tandem affinity purification (TAP) coupled to liquid chromatography-tandem mass spectrometry (LC-MS/MS). Our data revealed KAT2A (also known as GCN5) and KAT2B (also known as PCAF) as the top robust binding partners of MAGE-A10 (**Figure 2A and Table S1**). Both KAT2A and KAT2B are HAT enzymes share significant structural and functional similarities, including the presence of a conserved acetyltransferase domain that catalyzes the acetylation of lysine residues on histone tails, primarily on histone H3 at lysines 9 and 14 (H3K9 and H3K14), which are associated with transcriptional activation. We confirmed that MAGE-A10 interacts with KAT2A in H446 cells by co-immunoprecipitation (co-IP) (**Figure 2B**). Further analysis revealed that the MAGE-A10 directly binds KAT2A *in vitro* (**Figure 2C**).

**Figure 2.**
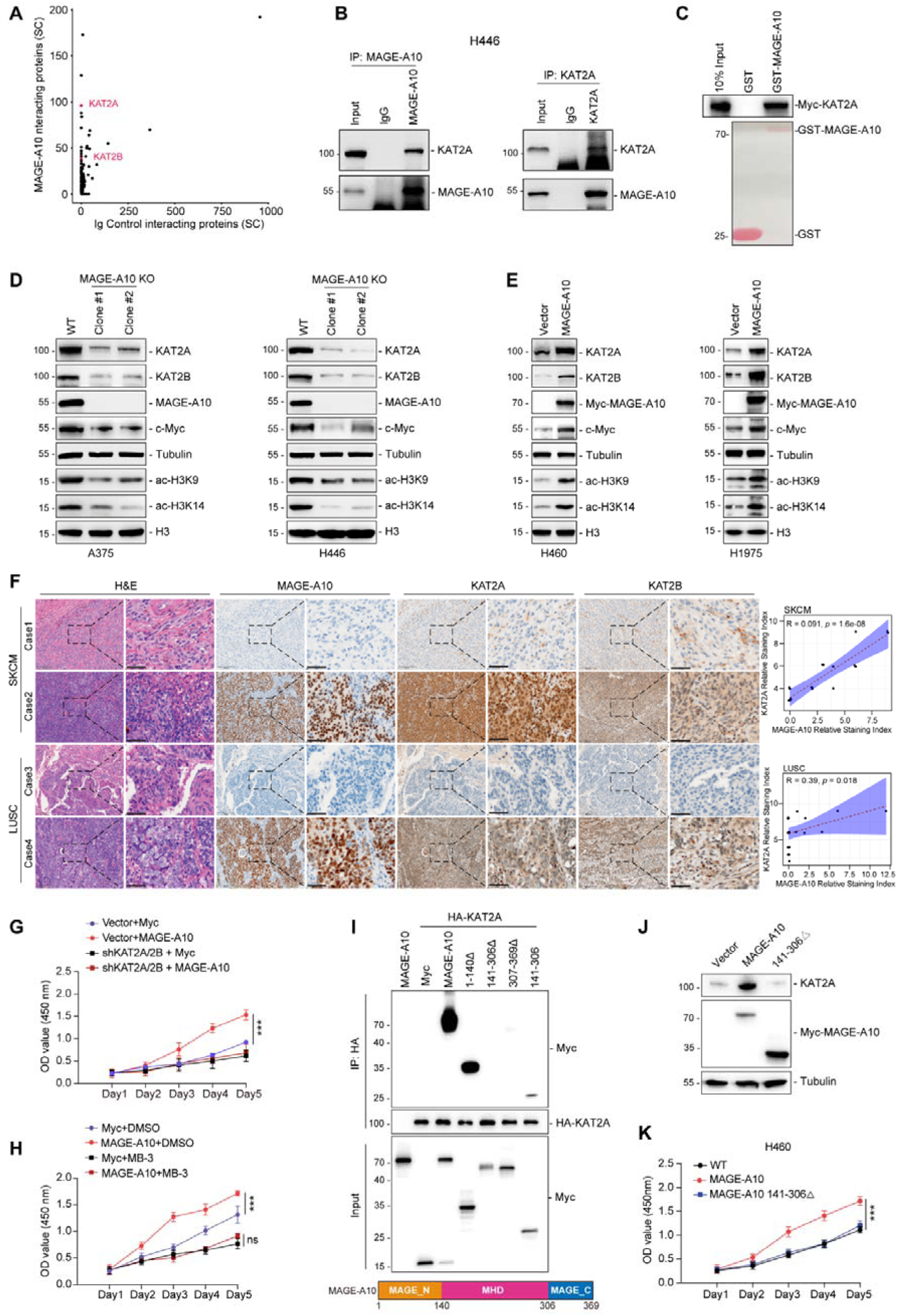
MAGE-A10 Promotes KAT2A/KAT2B Expression and Histone 3 Lysine Acetylation. (A) MAGE-A10 interacts with KAT2A/KAT2B proteins. HEK293 cells stably expressing TAP-vector or TAP-MAGE-A10 were subjected to pull-down followed by SDS-PAGE and LC-MS/MS. Spectral counts (SC) of the identified proteins in TAP-MAGE-A10 pull down, but absence in TAP-vector control, are shown. (B) MAGE-A10 interacts with KAT2A in cells. H446 cell lysates were subjected to immunoprecipitation (IP) with control IgG, anti-MAGE-A10, or anti-KAT2A antibodies. The immunoprecipitates were then blotted with the indicated antibodies. (C) Recombinant glutathione S-transferase (GST)-MAGE-A10, but not GST, binds *in vitro* translated Myc-KAT2A. (D) Knockout of MAGE-A10 decreases KAT2A/KAT2B protein levels and Histone 3 lysine acetylation. Wild-type or MAGE-A10 knockout A375 or H446 cells were blotted for the indicated proteins. (E) Over-expression of MAGE-A10 increases KAT2A/KAT2B protein levels and Histone 3 lysine acetylation. Indicated proteins were blotted from MYC-vector or MYC-MAGE-A10 over-expressing H460 and H1975 lung cancer cells, respectively. (F) Representative IHC images showing positive staining for MAGE-A10 in Skin Cutaneous Melanoma (SKCM) and Lung Squamous Cell Carcinoma (LUSC) tissues correlated with higher expression levels of KAT2A/KAT2B in the same patient samples. (G) Knockdown of KAT2A/KAT2B abolished MAGE-A10-induced cell proliferation advantage. KAT2A/KAT2B-knockdown H460 cells were transfected with indicated constructs. Cell proliferation was determined by CCK8 assay at indicated time points (n = 6 biological replicates). (H) Inhibition of KAT2A/KAT2B activity abolished MAGE-A10 induced cell proliferation. H460 cell proliferation assays were performed in control or MAGE-A10 overexpressing H460 cells with or without MB-3 treatment (n = 6 biological replicates). (I) Mapping the specific region of MAGE-A10 necessary for interaction with KAT2A. HEK293FT cells stably expressing HA-KAT2A were transfected with indicated MAGE-A10 constructs for 48 h before IP with anti-HA followed by SDS–PAGE and immunoblotting for anti-Myc. (J) MAGE-A10 141-306 mutant abolished the interaction with KAT2A and failed to regulate KAT2A expression. H460 cells were transfected with wild type or 141-306 deplete mutation MAGE-A10 for 48 h. Indicated proteins were blotted respectively. (K) MAGE-A10 141-306 mutant failed to promote cell proliferation. H460 cells were transfected with wild type or 141-306 deplete mutation MAGE-A10 for 48 h. Cell viability was determined by CCK8 assay at indicated time points (n = 6 biological replicates). Data are means ± SDs. Asterisks indicate significant differences from the control (*p<0.05, **p<0.01, ***p<0.001, n.s., not significant).

An intriguing mechanism of MAGE proteins is their ability to bind specific E3 ubiquitin ligases and modulate the ubiquitination of target proteins^37–39^. Based on this, we first examined whether the aberrant expression of MAGE-A10 regulates KAT2A and KAT2B protein levels. Surprisingly, knockout of MAGE-A10 resulted in decreased expression of both KAT2A and KAT2B, as well as reduced acetylation of their downstream targets H3K9 and H3K14, and decreased MYC expression in MAEG-A10 positive cancer cells (**Figure 2D** and **Figure S2A**). Conversely, introducing MAGE-A10 expression in MAGE-A10-negative cancer cell lines increased the protein levels of KAT2A and KAT2B, along with enhanced acetylation of H3K9 and H3K14, and elevated MYC expression (**Figure 2E** and **Figure S2B**). Furthermore, immunohistochemistry staining of SKCM and LUSC human specimens confirmed the presence of MAGE-A10 protein, which was accompanied by higher expression levels of KAT2A and KAT2B (**Figure 2F**). The positive correlation between MAGE-A10 and KAT2A/2B suggests that, unlike the known role of MAGE proteins as E3 adaptors promoting degradation, MAGE-A10 may regulate KAT2A/2B through an unknown mechanism.

To determine whether KAT2A/2B upregulation upon MAGE-A10 expression contributes to MAGE-A10-driven proliferation, we performed double knockdown of KAT2A/2B in H446 cells, with or without MAGE-A10 expression, and assessed cell proliferation rates. Overexpression of MAGE-A10 increased proliferation in wild-type H446 cells, but not in KAT2A/2B double knockdown cells (**Figure 2G**). Furthermore, inhibiting KAT2A activity with Butyrolactone 3 (MB-3) treatment also diminished the cell growth advantage in MAGE-A10 overexpressing cells (**Figure 2H**). To further determine whether MAGE-A10 regulation of cell growth depends on its ability to interact with KAT2A, we identified the region of MAGE-A10 required for binding KAT2A. Cell-based co-immunoprecipitation (co-IP) assays showed that the MAGE homology domain (MHD; amino acids 141-306) of MAGE-A10 is required for binding to KAT2A (**Figure 2I**). Deletion of the MHD in MAGE-A10 failed to increase the protein level of KAT2A (**Figure 2J**). Importantly, MHD-deleted MAGE-A10 could not induce cell growth in H460 cells compared to wild-type MAGE-A10 (**Figure 2K**). Collectively, these findings indicate that MAGE-A10-mediated cellular proliferation is dependent on its ability to bind KAT2A.

### MAGE-A10 Prevents KAT2A/KAT2B Autophagic Degradation via p62

To elucidate the mechanism by which MAGE-A10 increases KAT2A/2B protein levels, we first tested whether MAGE-A10 regulates KAT2A/2B mRNA transcription. Our results showed that neither knockdown nor overexpression of MAGE-A10 affected KAT2A/2B transcript levels (**Figure S3A** and **Figure S3B**). However, knockout of MAGE-A10 dramatically decreased the stability of KAT2A and KAT2B protein (**Figure 3A**). Consistently, overexpression of MAGE-A10 prolonged the half-life of the KAT2A and KAT2B protein (**Figure 3B**). These results indicated MAGE-A10 prevents KAT2A and KAT2B protein degradation.

**Figure 3.**
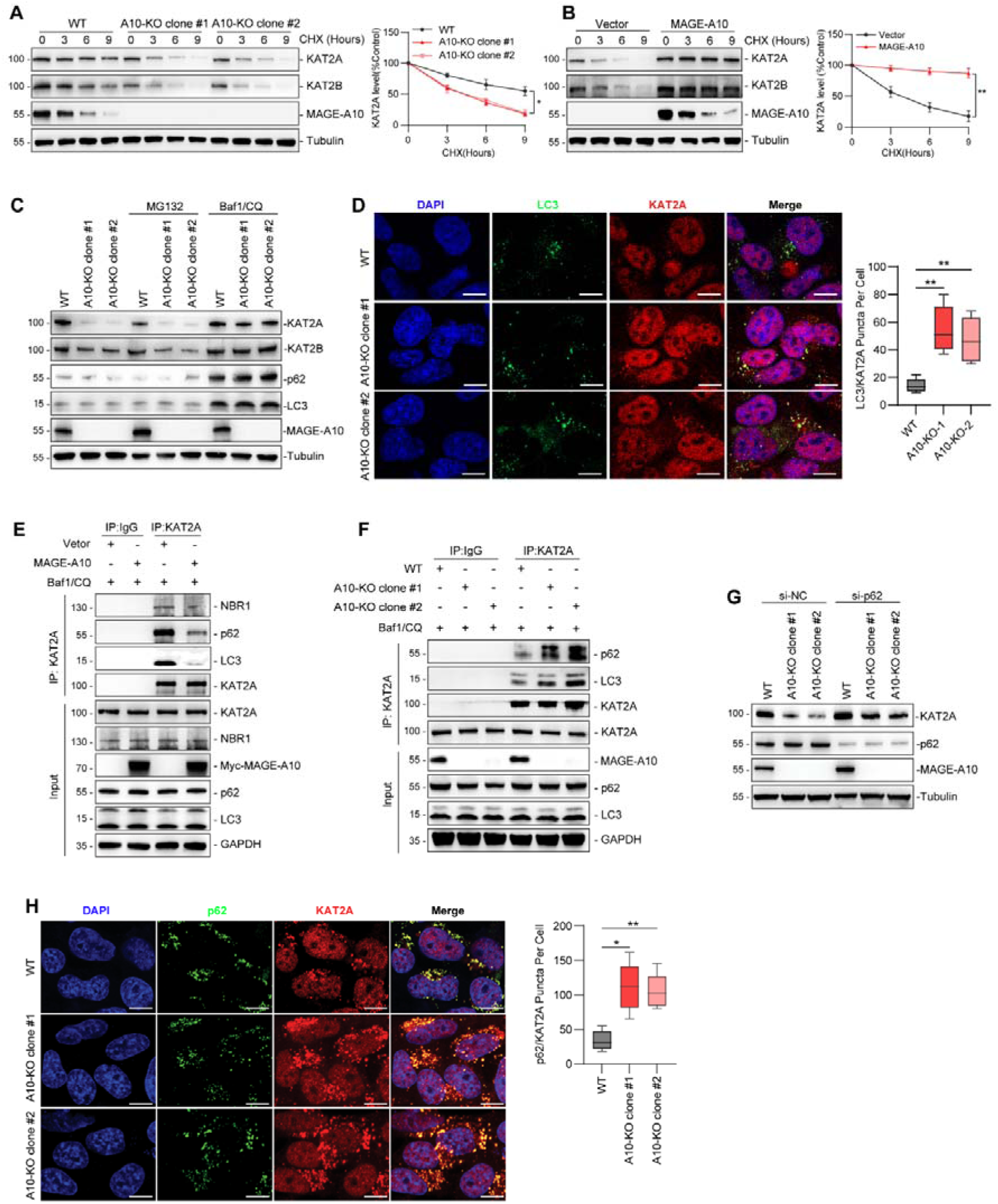
MAGE-A10 Prevents KAT2A/KAT2B Autophagic Degradation via p62. (A) Knockout of MAGE-A10 decreased KAT2A/KAT2B protein stability in A375 cells. MAGE-A10 wild-type or knockout A375 cells were treated with 100 mg/mL cycloheximide for the indicated times. Cell lysates were immunoblotted and quantitated (n = 3). (B) Over-expression of MAGE-A10 increased KAT2A/KAT2B protein stability in H460 cells. MAGE-A10 was stably expressed in H460 cells and were treated with 100 mg/mL cycloheximide for the indicated times. Cell lysates were immunoblotted and quantitated (n = 3). (C) Knockout MAGE-A10 promoted KAT2A degradation via autophagy. MAGE-A10 wild-type or knockout A375 cells were treated with 10 mM MG132 or Baf1 and CQ for 4 h before immunoblotting. (D) Knockout of MAGE-A10 enhanced the co-localization of KAT2A with LC3 during autophagy. Wild-type and MAGE-A10 knockout A375 cells were treated with 200 nM Baf1 and 50 μM CQ for 4 hours. Endogenous LC3 and KAT2A were detected using anti-LC3 and anti-KAT2A antibodies, respectively. The number of KAT2A/LC3 co-localized fluorescent puncta per cell was quantified in 40 cells for each data point. Scale bar = 10 μm. (E) MAGE-A10 inhibited p62 or LC3 binding to KAT2A. H460 cells stably expressing MYC-vector or MYC-MAGE-A10 were treated with 200 nM Baf1 and 50 μM CQ for 4 h followed by IP with anti-KAT2A antibodies. The indicated proteins were immunoblotted. (F) Depletion of MAGE-A10 enhanced p62 or LC3 binding to KAT2A. MAGE-A10 Wild-type or knockout A375 stable cell lines were treated with 200 nM Baf1 and 50 μM CQ for 4 h followed by IP with anti-KAT2A antibodies. The indicated proteins were immunoblotted. (G) MAGE-A10 regulated KAT2A degradation mediated by p62. Wild type or MAGE-A10 knock out cells were transfected by control- or p62-targeting siRNA for 72 h, respectively. Cell lysates were immunoblotted by indicated proteins. (H) Knockout of MAGE-A10 increased the co-localization of KAT2A with p62 during autophagy. Cells were treated in the same manner as described for Figure (D) and subsequently analyzed via immunofluorescence. The number of KAT2A/p62 co-localized fluorescent puncta per cell was quantified in 40 cells for each data point. Scale bar = 10 μm. Data are means ± SDs. Asterisks indicate significant differences from the control (*p<0.05, **p<0.01, ***p<0.001, n.s., not significant).

Proteasome and autophagy are two primary pathways for protein degradation in cells^40,41^. The pathway mediating the degradation of KAT2A/2B remains unclear. Previous studies by Changhan Ouyang *et al.* have shown that KAT2A undergoes autophagic degradation in human aortic smooth muscle cells^42^. In contrast, other groups have provided evidence that KAT2A is ubiquitinated *in vivo* and degraded *via* the proteasome pathway^43,44^. To evaluate the pathway by which MAGE-A10 mediates KAT2A degradation in cancer cells, we treated cells with the proteasome inhibitor MG132 or the lysosome inhibitors chloroquine (CQ) and Bafilomycin A1 (BafA1). Our results showed that CQ/BafA1 treatment resulted in increased levels of LC3-II and SQSTM1 (p62) proteins, along with an increase in KAT2A/2B protein levels, indicating that defective autophagy reduces KAT2A/2B protein degradation. Conversely, proteasome inhibition with MG132 had little effect on KAT2A/2B protein levels (**Figure 3C**). Consistently, overexpressing MAGE-A10 no longer increased the protein levels of KAT2A in CQ/BafA1-treated cells (**Figure S3C**). Overall, our results indicate that MAGE-A10 prevents KAT2A from undergoing autophagic degradation.

To further validate that MAGE-A10 inhibits the autophagic degradation of KAT2A, we examined the localization of KAT2A in autophagosomes using immunostaining in cells. Fluorescence microscopy analysis revealed that KAT2A is mainly localized in nuclei. However, activating autophagy induced KAT2A transport into cytoplasm (**Figure S3D**). To validate whether cytoplasm transported KAT2A prone to autophagic degradation, we assessed the co-localization of KAT2A with LC3. Our results showed that KAT2A co-localized with LC3 puncta in the cytoplasm. Moreover, knockout of MAGE-A10 increased the co-localization of KAT2A with LC3 (**Figure 3D**). Conversely, introducing MAGE-A10 into MAGE-A10-negative cells prevented the co-localization of KAT2A with LC3 (**Figure S3E**). These findings confirm that MAGE-A10 plays a crucial role in preventing the targeting of KAT2A to autophagic degradation in the cytoplasm.

Given that KAT2A is primarily localized in the nuclei but is transported into the cytoplasm and co-localizes with LC3 for autophagic degradation, it suggests that KAT2A degradation may occur through selective autophagy. Selective autophagy is mediated by receptors such as p62 (SQSTM1) or NBR1 (Neighbor of BRCA1 gene 1) that recognize and bind specific substrates. Both p62 and NBR1 contain LC3-interacting regions (LIRs) and ubiquitin-binding domains (UBDs), enabling them to act as bridges between LC3 proteins and ubiquitinated substrates^45,46^. To elucidate the specific adaptors involved in the selective autophagic degradation of KAT2A, we treated H460 cells with CQ/BafA1 to block lysosomal activity and performed immunoprecipitation (IP) of endogenous KAT2A to assess its interactions with autophagy receptors. Our results demonstrate a strong interaction between KAT2A and p62. Importantly, MAGE-A10 prevents the interaction between KAT2A and p62, as well as the subsequent binding of KAT2A to LC3 (**Figure 3E**). Conversely, knockout of MAGE-A10 in cells enhanced the interaction between KAT2A and p62, along with the binding of KAT2A to LC3 (**Figure 3F**). Additionally, knockdown of p62 significantly rescued KAT2A protein levels in MAGE-A10 knockout cells (**Figure 3G**). As expected, KAT2A co-localized with p62 during autophagy, and this co-localization was significantly enhanced in MAGE-A10 knockout cells (**Figure 3H**). Moreover, expression of MAGE-A10 prevented the co-localization of KAT2A with p62 (**Figure S3F**). These findings indicate that MAGE-A10 inhibits the selective autophagic degradation of KAT2A by preventing its interaction with p62.

### MAGE-A10 Inhibits K63-linked Polyubiquitination of KAT2A Mediated by the CUL4A E3 Ubiquitin Ligase

Considering that p62 mediates selective autophagy by recognizing ubiquitinated proteins, we next investigated the ubiquitination status of KAT2A. Our analyses revealed that MAGE-A10 expression decreases the ubiquitination of KAT2A (**Figure 4A**). Conversely, knockout of MAGE-A10 significantly increased KAT2A ubiquitination (**Figure 4B** and **Figure S4A**), suggesting that MAGE-A10 regulates KAT2A ubiquitination in cells. To further elucidate the type of ubiquitin linkage involved, we examined the ubiquitination of KAT2A and found that it predominantly involves K63-specific chains (**Figure 4C** and **Figure S4B**). Additionally, perturbation of MAGE-A10 expression primarily affected the formation of K63-linked ubiquitin chains on KAT2A (**Figure 4D** and **Figure 4E**). These results suggest that MAGE-A10 specifically regulates the K63-linked ubiquitination of KAT2A.

**Figure 4.**
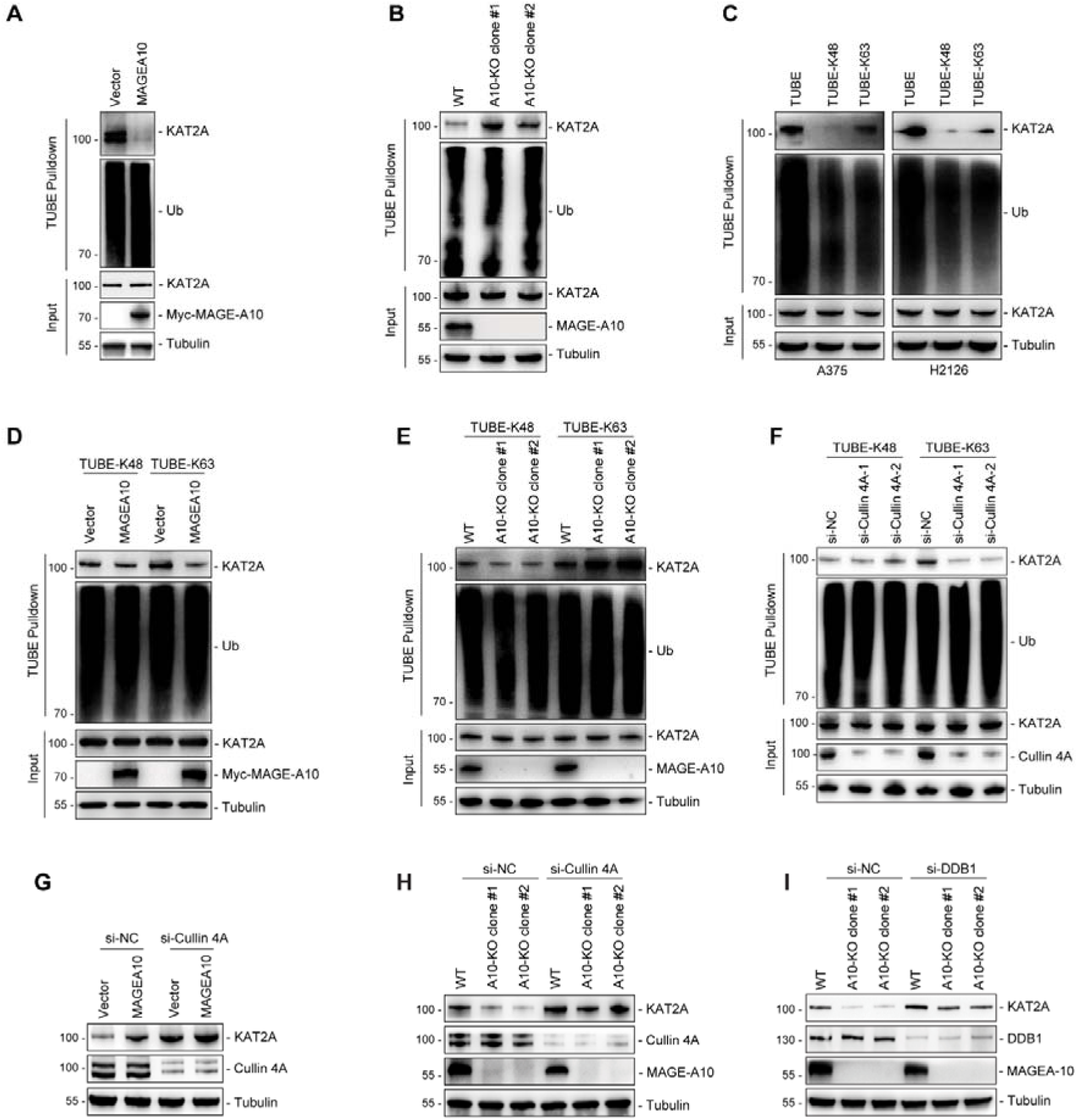
MAGE-A10 Inhibits K63-linked Polyubiquitination of KAT2A Mediated by the CUL4A-DDB1 E3 Ubiquitin Ligase Complex. (A) Expression of MAGE-A10 inhibited KAT2A ubiquitination. Ubiquitinated proteins from MYC-vector or MYC-MAGE-A10 stably expressing H460 cells were treated with 200 nM Baf1 and 50 μM CQ for 4 h, then cells were isolated with tandem ubiquitin binding entity (TUBE)-agarose followed by SDS-PAGE and immunoblotting for endogenous KAT2A. (B) Knockout of MAGE-A10 increased KAT2A ubiquitination. MAGE-A10 wild-type or knockout A375 cells were treated and isolated as described in Figure A and subsequently immunoblotting for indicated proteins. (C) Ubiquitin chain type on KAT2A. A375 and H2126 cells were treated as described in Figure A and subsequently isolated using universal, K48-specific, or K63-specific TUBE-agarose, respectively. Endogenous KAT2A levels were then determined by immunoblotting. (D) Expression of MAGE-A10 inhibited K63-linked polyubiquitination of KAT2A. Ubiquitinated proteins from MYC-vector or MYC-MAGE-A10 stably expressing H460 cells were treated with 200 nM Baf1 and 50 μM CQ for 4 h, then cells were isolated with K48-specific or K63-specific TUBE-agarose followed by SDS-PAGE and immunoblotting for endogenous KAT2A. (E) Knockout of MAGE-A10 increased K63-linked polyubiquitination of KAT2A. MAGE-A10 wild-type or knockout A375 cells were treated and isolated as described in Figure D and subsequently immunoblotting for indicated proteins. (F) CUL4A induces K63-linked polyubiquitination of KAT2A. Ubiquitinated proteins from MAGE-A10 wild-type or knockout A375 cells were transfected with si-NC or si-Cullin 4A for 72 hours for 72 h, then cells were isolated with K48-specific or K63-specific TUBE-agarose followed by SDS-PAGE and immunoblotting for endogenous KAT2A. (G) Knockdown of CUL4A rescued KAT2A protein stability in MAGE-A10 knockout cells. MYC-vector or MYC-MAGE-A10 stably expressing H460 cells were treated with si-NC or siCUL4A. Cell lysates were immunoblotted for the indicated proteins. (H) Knockdown of CUL4A rescued KAT2A protein stability in MAGE-A10 knockout cells. Wild-type or MAGE-A10 knockout A375 cells were treated with si-NC or siCUL4A. Cell lysates were immunoblotted for the indicated proteins. (I) Knockdown of DDB1 abolished the regulation of KAT2A degradation by MAGE-A10. Wild-type or MAGE-A10 knockout A375 cells were treated with si-NC or siDDB1. Cell lysates were immunoblotted for the indicated proteins.

Previous studies have suggested that the E3 ubiquitin ligase CUL4A complex mediates the ubiquitination of KAT2A^44,47^. As one member of the evolutionarily conserved Cullin-RING E3 family proteins, the CRL4 E3 ligase complex is composed of a scaffold protein Cullin4 (CUL4), and the adaptor protein DNA damage-binding protein 1 (DDB1)^48–50^. To investigate whether the CUL4A complex is involved in the regulation of KAT2A turnover by MAGE-A10, CUL4A was knocked down in both wild-type and MAGE-A10 overexpressing cell lines. Knockdown of CUL4A resulted in decreased K63-linked polyubiquitination of KAT2A (**Figure 4F**). Furthermore, the prevention of KAT2A degradation by MAGE-A10 was shown to be dependent on CUL4A and DDB1 (**Figure 4G-I**). These findings indicate that MAGE-A10 modulates KAT2A stability via a CUL4A-DDB1-dependent mechanism.

### Regulation of KAT2A Ubiquitination Is Essential for MAGE-A10-Induced Tumorigenesis

Next, to test whether CUL4A mediates the regulation of KAT2A ubiquitination by MAGE-A10, K63-linked ubiquitination of KAT2A was assessed in CUL4A wild-type and knockdown cell lines with or without MAGE-A10 expression. As shown in Figure 5A, MAGE-A10 expression decreased K63-linked ubiquitination of KAT2A. In contrast, the knockdown of CUL4A decreased the ubiquitination of KAT2A and abolished the regulation of KAT2A ubiquitination by MAGE-A10 (**Figure 5A**). To further confirm this observation, an *in vitro* ubiquitination assay was performed. Our results showed that CUL4A complexes were able to ubiquitinate KAT2A *in vitro*; however, the addition of purified MAGE-A10 reduced the ubiquitination of KAT2A (**Figure 5B**). Taken together, our data demonstrate that MAGE-A10 prevents KAT2A ubiquitination by the CUL4A E3 complex both *in vivo* and *in vitro*.

**Figure 5.**
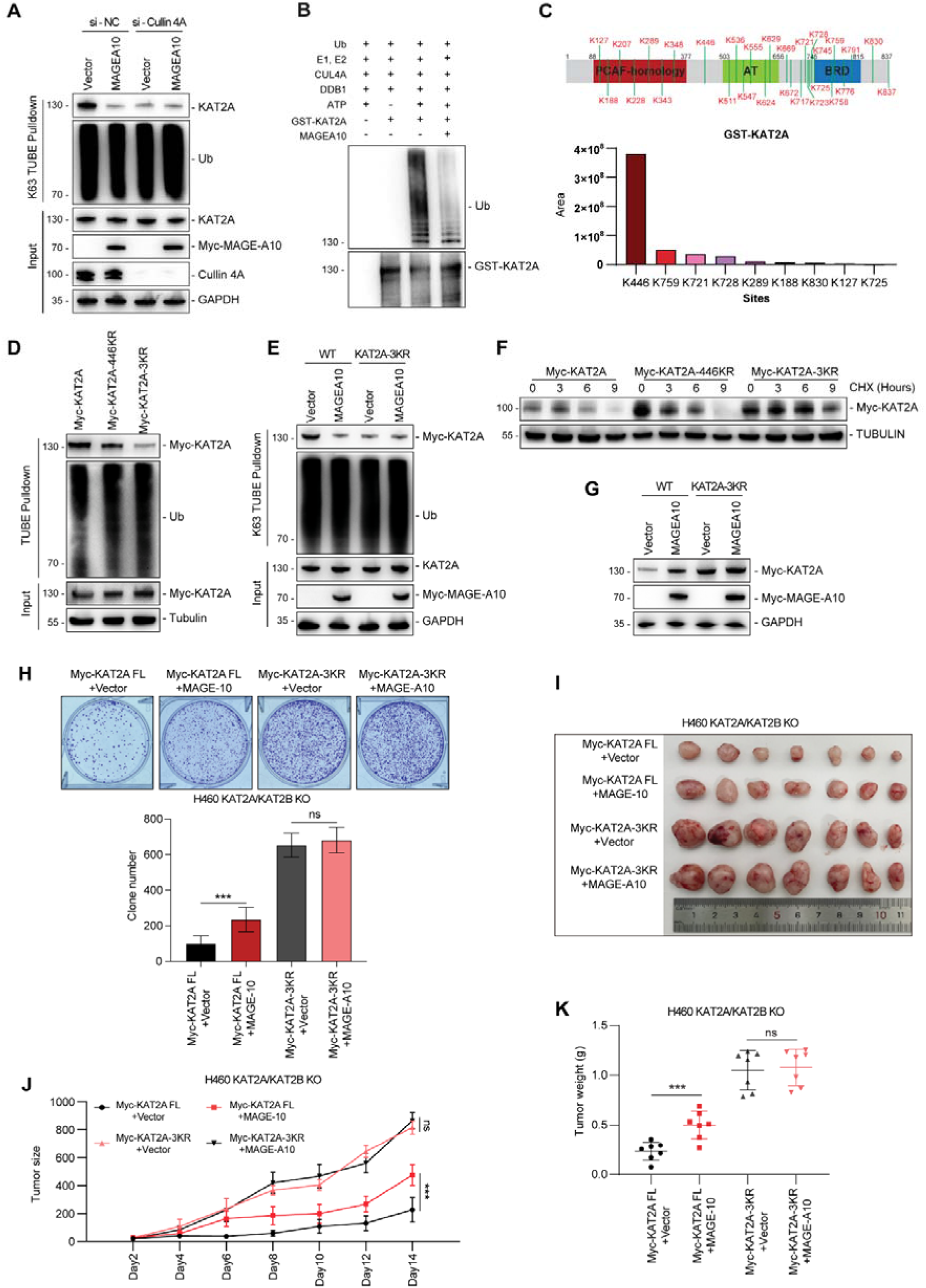
Regulation of KAT2A Ubiquitination Is Essential for MAGE-A10-Induced Tumorigenesis. (A) Expression of MAGE-A10 inhibited K63-linked polyubiquitination of KAT2A through CUL4A. Ubiquitinated proteins from MYC-vector or MYC-MAGE-A10 stably expressing H460 cells were treated with si-NC or si-CUL4A for 72h, then cells were isolated with K63-specific TUBE-agarose followed by SDS-PAGE and immunoblotting for endogenous KAT2A. (B) MAGE-A10 prevented KAT2A ubiquitination by CUL4A-DDB1 *in vitro*. Incubation of immunopurified CUL4A-DDB1 and KAT2A in an *in vitro* ubiquitin ligase assay increased the formation of KAT2A ubiquitinated species. The addition of MAGE-A10 suppressed the KAT2A ubiquitination by CUL4A-DDB1. (C) Identification of KAT2A ubiquitination sites using an *in vitro* ubiquitin ligase assay. All lysine residues of KAT2A are depicted on the domain map (top). Lysine residues identified as ubiquitinated by mass spectrometry (MS) are highlighted with frames. The quantification of these ubiquitinated lysine sites is presented in the bar plot at the bottom. (D) Analysis of KAT2A ubiquitination following site-directed mutagenesis. Ubiquitinated proteins from wild-type (WT) KAT2A and KAT2A mutants with three lysine residues replaced by alanine (K to A) were used to assess their role in ubiquitination. A375 cells were treated and then cells were isolated with TUBE-agarose followed by SDS-PAGE and immunoblotting for endogenous Myc-KAT2A. (E) MAGE-A10 did not regulate KAT2A ubiquitination after site-directed mutagenesis. wild-type (WT) KAT2A and KAT2A mutants with three lysine residues replaced by alanine (K to A) cells were transfected with Myc-Vector or Myc-MAGE-A10 for 48h and then then cells were isolated with TUBE-agarose followed by SDS-PAGE and immunoblotting for endogenous Myc-KAT2A. (F) Assessment of KAT2A stability following site-directed mutagenesis. The impact of lysine-to-alanine mutations on KAT2A protein degradation was evaluated by using 100 mg/mL cycloheximide treatment for the indicated times. Cell lysates were immunoblotted for KAT2A protein. (G) MAGE-A10 no longer increased the protein levels of the 3KR mutant. (H) Mutation in the ubiquitination sites of KAT2A enhances cell growth and abolishes KAT2A-induced regulation of cell growth *in vitro*. Soft agar experiments were performed to access cell growth of KAT2A WT and mutants with or without MAGE-A10 expression (n = 6 per group). Data are mean ± SD, ***p < 0.001. (I-K) Stable expression of MAGE-A10 in KAT2A ubiquitination sites mutation does not increase xenograft tumor growth in mice (n = 7 per group). Data are mean ± SD, ***p < 0.001.

We further mapped the ubiquitination sites of KAT2A that are regulated by MAGE-A10. Eight lysine residues were identified as potential ubiquitination sites by mass spectrometry from the *in vitro* ubiquitination assay (**Table S2**). Among these, K446 at the N-terminal, and K759, K721, and K728 at the C-terminal exhibited the highest levels of ubiquitination on KAT2A (**Figure 5C**). Follow-up analysis on mutants of the potential KAT2A ubiquitination sites (KR mutants) revealed that K446 mutation had little effect on the ubiquitination, while the three K-to-R mutants on K759, K721, and K728 (3KR) almost abolished the ubiquitination on KAT2A. These results indicate K63 ubiquitination of KAT2A predominantly occurs at K721, K728, and K759 (3KR) (**Figure 5D**). We further tested whether the ubiquitination at these sites is regulated by MAGE-A10. As shown in Figures 5E, overexpression of MAGE-A10 decreased the ubiquitination of wild-type (WT) KAT2A; however, overexpression of MAGE-A10 did not affect the ubiquitination levels of the KAT2A 3KR mutant (**Figures 5E**). These results suggest that these sites are the major ubiquitination sites regulated by MAGE-A10. To investigate the functional significance of KAT2A ubiquitination, we first examined the protein half-life of WT KAT2A and the 3KR mutant. Indeed, the 3KR mutant exhibited a significantly longer half-life compared to WT KAT2A (**Figure 5F**). Furthermore, MAGE-A10 no longer increased the protein levels of the 3KR mutant (**Figure 5G**). These findings indicate that MAGE-A10 regulates the stability of KAT2A through specific ubiquitination sites.

To study the impact of these KAT2A ubiquitination site mutations on tumor growth, we reconstituted KAT2A knockout H460 cells with either WT KAT2A or the 3KR mutant and examined tumor growth both *in vitro* and *in vivo*. Mutation of the KAT2A ubiquitination sites increased colony formation and abolished the growth advantage conferred by MAGE-A10 expression (**Figure 5H**). Xenograft experiments further demonstrated that the KAT2A mutations significantly enhanced tumor growth and diminished the regulatory effect of MAGE-A10 on KAT2A (**Figure 5I-5K**). These findings highlight the importance of KAT2A ubiquitination regulation in MAGE-A10-induced tumor growth.

### MAGE-A10 Antagonizes DDB1 for the Association with KAT2A

In combination with CUL4A, DDB1 has been proposed to either directly dock a substrate to the E3 machinery or indirectly recruit a substrate through additional adaptor proteins that specifically recognize the target protein^48–50^. Essentially, DDB1 acts as a flexible platform to bring the substrate and the E3 complex together, depending on the cellular context. A previous study has indicated that DTL may serve as a substrate receptor protein for the CUL4A-DDB1 complex, mediating KAT2A degradation^44^. To explore the mechanism by which MAGE-A10 prevents the ubiquitination of KAT2A by CUL4A, we first examined the partners involved in the E3 complex. Indeed, our results showed that DDB1, CUL4A, and DTL all bind with endogenous KAT2A in cells (**Figure 6A**). However, only DDB1 demonstrated direct binding with purified GST-KAT2A *in vitro* (**Figure 6B-D**). Given that both CUL4A and DDB1 interact with KAT2A, it is possible that MAGE-A10 may compete with DDB1 for binding to KAT2A. To explore this possibility, the interaction between KAT2A and DDB1 was examined by introducing MAGE-A10 expression in MAGE-A10-negative H460 and M8 cells. As expected, expression of MAGE-A10 decreased the interactions between KAT2A with DDB1 and CUL4A (**Figure 6E** and **Figure S5A**). Consistently, knockout of MAGE-A10 in MAGE-A10-positive H446 and A375 cells enhanced the interaction between KAT2A and DDB1 (**Figure 6F** and **Figure S5B**). These findings suggest that MAGE-A10 disrupts the interaction between KAT2A and the CUL4A-DDB1 complex, thereby preventing KAT2A ubiquitination.

**Figure 6.**
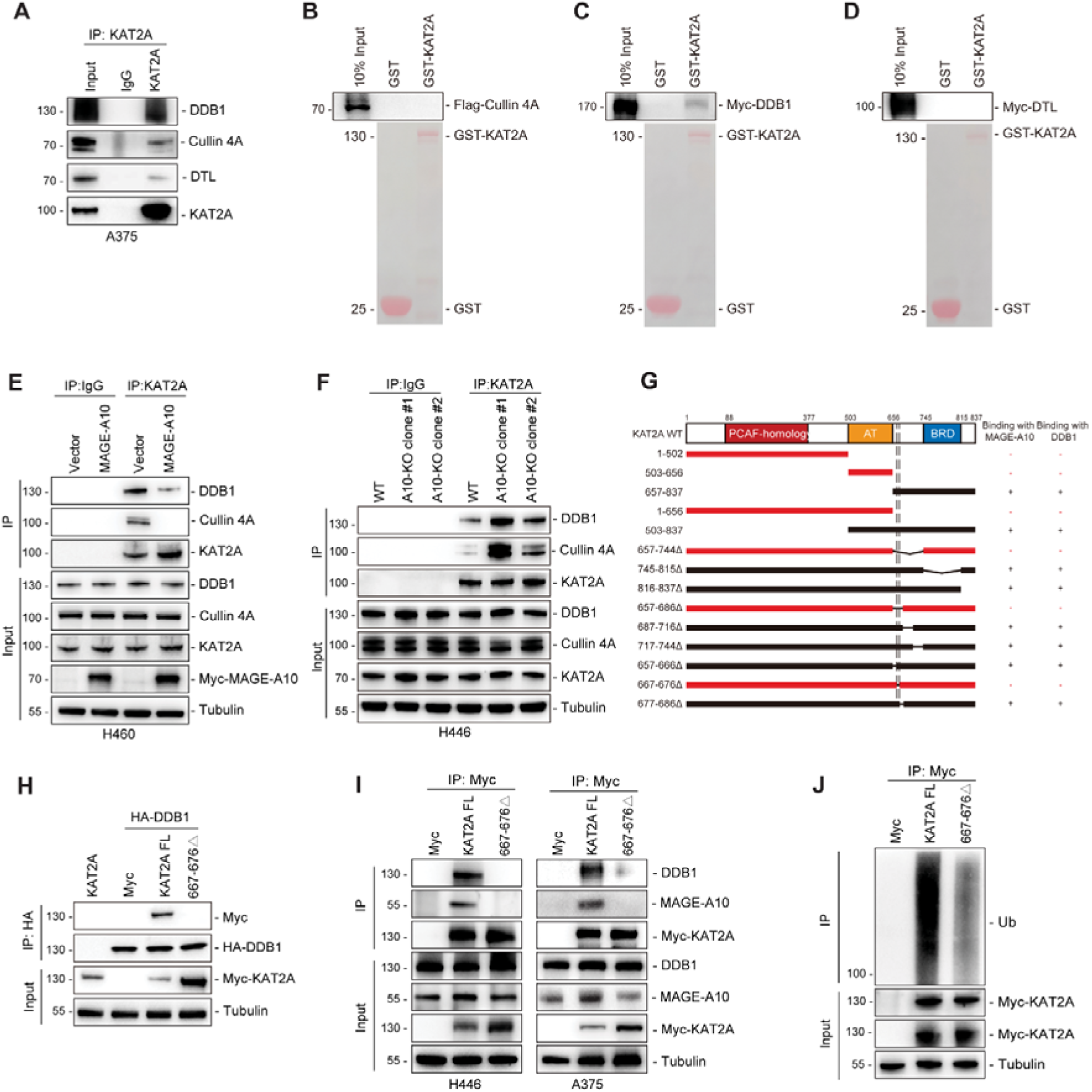
MAGE-A10 antagonizes DDB1 for the association with KAT2A. (A) KAT2A interacts with DDB1 *in vivo*. Endogenous KAT2A protein was immunoprecipitated from the A375 cell line and subsequently analyzed by immunoblotting for the indicated proteins. (B-D) KAT2A directly interacts with DDB1 *in vitro* as demonstrated by GST pull-down assays. Recombinant GST-KAT2A was incubated with in vitro translated Flag-CUL4A (B), Myc-DDB1 (C), or Myc-DTL (D). The complexes were pulled down using glutathione sepharose beads, eluted, separated by SDS-PAGE, and analyzed by immunoblotting. (E) MAGE-A10 represses the interaction between KAT2A and the CUL4A-DDB1 complex. Endogenous KAT2A protein was immunoprecipitated from H460 wild-type (WT) or MAGE-A10 overexpressing stable cell lines. The interaction between KAT2A and the CUL4A-DDB1 complex was assessed by immunoblotting. (F) Deletion of MAGE-A10 increases the interaction between KAT2A and the CUL4A-DDB1 complex. Endogenous KAT2A protein was immunoprecipitated from either H446 wild-type (WT) or with MAGE-A10 knockout stable cell lines. The interaction between KAT2A and the CUL4A-DDB1 complex was examined through immunoblotting. (G) Identification of binding motif on KAT2A for DDB1 and MAGE-A10 interaction. HEK293FT cells stably expressing HA-DDB1 were transfected with the indicated constructs for 48 h before IP with anti-HA followed by SDS-PAGE and immunoblotting for anti-Myc. (H) Mutation of the KAT2A binding sites abolishes interactions with DDB1. A375 cell lines were transfected with Myc-KAT2A, KAT2A 667-676 mutation and HA-DDB1 for 48 h before IP with anti-HA followed by SDS-PAGE and immunoblotting for indicated proteins. (I) Mutation of the KAT2A binding sites abolishes interactions with both MAGE-A10 and DDB1. H446 cell lines were transfected with Myc-KAT2A or KAT2A 667-676 mutation for 48 h before IP with anti-Myc followed by SDS-PAGE and immunoblotting for indicated proteins. (J) Mutations in the KAT2A binding sites for DDB1 decrease KAT2A ubiquitination. A375 cell lines were transfected with Myc-KAT2A or KAT2A 667-676 mutation for 48 h before IP with anti-Myc followed by SDS-PAGE and immunoblotting for ubiquitin.

To further investigate whether MAGE-A10 competes with DDB1 for binding to the same region on KAT2A, we mapped the interaction regions on KAT2A (**Figure S6A-E**). Interestingly, amino acid 667-676 on C-terminal of KAT2A represent the same binding region for both MAGE-A10 and DDB1 (**Figure 6G-H and S6F-I**). Depletion of 667-676 (667-676Δ) abolished the interaction of KAT2A with both MAGE-A10 and DDB1 in H446 and A375 cells (**Figure 6I**). More importantly, compared with wild type KAT2A, mutation of 667-676 decreased the ubiquitination of KAT2A (**Figure 6J**). Overall, our findings suggest that MAGE-A10 and DDB1 compete for the same binding site on KAT2A, influencing its ubiquitination and stability.

### KAT2A in turn Promotes the Transcription of MAGE-A10 and Forms a Positive Feedback Loop

The aberrant expression of MAGE genes, including MAGE-A10, is a common feature in various cancers^28–30^. Despite extensive research, the underlying mechanisms driving this aberrant expression remain largely mysterious. One intriguing question is which factors regulate the expression of MAGE genes in cancer cells. Given that KAT2A is known to mediate histone acetylation, we hypothesized that KAT2A might influence the expression of MAGE-A10. Surprisingly, increasing expression of KAT2A enhanced the protein levels of MAGE-A10 (**Figure 7A**), while knockout of KAT2A decreased the expression of MAGE-A10 (**Figure 7B**). Furthermore, we investigated the impact of proteasomal degradation on MAGE-A10 protein levels in KAT2A-deficient cells. Treatment with proteasome inhibition MG132 had little effect on MAGE-A10 protein levels in KAT2A knockout cells (**Figure 7C**). This finding, along with the observation that KAT2A knockout did not affect the stability of MAGE-A10 protein (**Figure 7D**), suggests that KAT2A may regulate MAGE-A10 expression at the transcriptional level rather than through protein stability mechanisms. Consistent with this hypothesis, we found that increased expression of KAT2A promoted the transcriptional activation of *MAGE-A10* (**Figure 7E**), while KAT2A knockout reduced *MAGE-A10* gene expression (**Figure 7F**). Collectively, our results provided compelling evidence that KAT2A plays a crucial role in the transcriptional regulation of *MAGE-A10*, highlighting a novel epigenetic mechanism in the control of MAGE-A10 expression.

**Figure 7.**
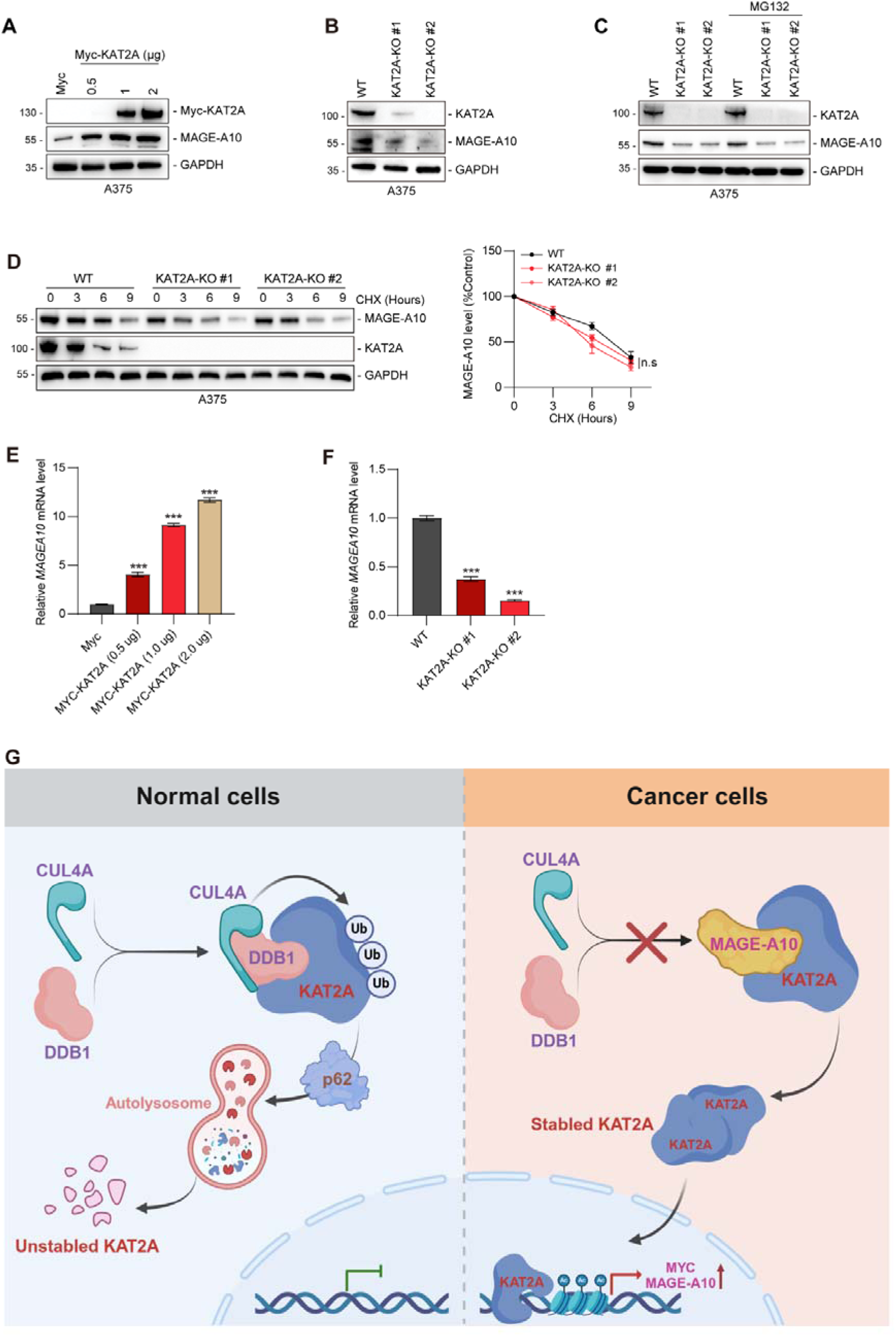
KAT2A in turn promotes the transcription of MAGE-A10 and forms a positive feedback loop. (A) Overexpression of KAT2A increased the expression of MAGE-A10. A375 cells were transfected with either control or increasing amount of Myc-KAT2A plasmid for 72 hours. Cells were then lysed and immunoblotted for the indicated proteins. (B) Knockout of KAT2A decreased the expression of MAGE-A10. WT or MAGE-A10 knockout A375 cells were lysed and immunoblotted for the indicated proteins. (C) MG132 treatment did not rescue MAGE-A10 expression in KAT2A knockout cells. KAT2A wild-type or KAT2A knockout A375 cell lines were treated with 10 μM MG132 for 6 hours. KAT2A and MAGE-A10 protein expressions were examined by immunoblotting. (D) KAT2A showed no effect on the stability of MAGE-A10. KAT2A wild-type or KAT2A knockout A375 cells were treated with 100 mg/mL cycloheximide for the indicated times. Cell lysates were immunoblotted and quantitated. (E) Overexpression of KAT2A increases MAGE-A10 mRNA transcription levels. A375 cells were transfected with increasing amounts of KAT2A, and MAGE-A10 mRNA levels were measured by qPCR (n=3). (F) Knockout of KAT2A decreases MAGE-A10 mRNA transcription. MAGE-A10 mRNA levels were determined by qPCR in KAT2A wild-type and knockout A375 stable cell lines (n=3). (G) Model for the MAGE-A10-KAT2A positive regulation feedback loop. MAGE-A10 antagonizes the association of CUL4A-DDB1 with KAT2A at the same binding site, thereby preventing KAT2A protein degradation through p62-mediated autophagy. Stabilized KAT2A in turn boosts the transcription of MAGE-A10 and forms a positive feedback loop. Data are means ± SDs. Asterisks indicate significant differences from the control (*p<0.05, **p<0.01, ***p<0.001, n.s., not significant).

In summary, our study has unveiled a novel mechanism by which the cancer-testis antigen MAGEA10 plays a role in orchestration of oncogenic processes. We discovered that MAGEA10 prevents KAT2A from degradation through the autophagy-lysosome pathway by blocking the interaction between KAT2A and DDB1. The consequent stabilization of KAT2A further amplifies MAGEA10 transcription, forming a positive feedback loop that activates the initiation and progression of tumors (**Figure 7G**).

## DISCUSSION

KAT2A and KAT2B are crucial histone acetyltransferases that play significant roles in gene regulation by acetylating histones and non-histone proteins^14,51,52^. These enzymes are pivotal in various biological processes, including DNA repair, cell cycle progression, and apoptosis^9–13^. Dysregulation of KAT2A and KAT2B has been implicated in several diseases, particularly cancer, where aberrant histone acetylation can lead to uncontrolled gene expression and tumorigenesis^17–23^. Despite their importance, the precise mechanisms regulating the stability and activity of KAT2A and KAT2B in cancer remain poorly understood. In this study, we identify MAGE-A10 as a novel cancer-specific regulator that stabilizes KAT2A and KAT2B, thereby enhancing their functional activity and contributing to oncogenesis.

MAGE-A10 is mainly expressed in the testis but is frequently turned on in many tumor types, including lung, melanoma, and bladder cancer^28–31^. Our findings suggest that activation of MAGE-A10 in cancer is not simply a passenger event during cellular transformation and tumorigenesis, but rather MAGE-A10 is a driver gene that supports multiple phenotypes associated with tumorigenesis. We propose that a critical oncogenic function of MAGE-A10 is the upregulation of KAT2A/2B protein expression, which disrupts the homeostasis of histone acetylation and consequently boosts the expression of oncogenes such as MYC. Our study demonstrated that MAGE-A10 expression promotes tumor cell proliferation in a KAT2A-dependent manner. In line with this, we showed that MAGE-A10 expression correlates with higher expression of KAT2A in patients. Consistently, patients with high levels of MAGE-A10 expression have a poorer prognosis^29,53,54^. Therefore, MAGE-A10 may serve as an important biomarker for predicting worse prognosis in cancer patients.

In recent years, the role of KAT2A/2B in cancer biology has become increasingly apparent, with growing evidence demonstrating its oncogenic function^14–16^. The initial connection of KAT2A/2B to cancer was established when these enzymes were found to acetylate and activate key oncogenic transcription factors, such as c-Myc and E2F^24,25^. However, KAT2A/2B exerts pleiotropic effects on cells by acetylating a variety of substrates, including histone H3, p53, and NF-κB, thereby altering cellular metabolic processes^55–57^. Future studies investigating whether MAGE-A10 influences KAT2A/2B acetylation of non-histone targets will be valuable. Regardless, our study indicates that MAGE-A10 drives tumorigenesis by modulating KAT2A/2B. Interestingly, KAT2A/2B inhibitors, either alone or in combination with other treatments, have been shown to reduce malignant growth^58^. Thus, MAGE-A10 expression status may serve as a useful biomarker for selecting patients with the greatest potential response to KAT2A/2B inhibitors.

How does MAGE-A10 promote the expression of KAT2A/2B? Previous studies indicate that specific E3 ubiquitin ligases may target KAT2A for proteasomal degradation, implicating the ubiquitin-proteasome system in its turnover^43,44^. Concurrently, other studies suggest that autophagy-related proteins may be involved in the degradation of KAT2A, pointing to a potential role for autophagic pathways^42,59,60^. Nevertheless, the precise molecular players and regulatory signals governing KAT2A stability remain incompletely elucidated. We found that KAT2A degradation occurs mainly through p62-mediated selective autophagy. Mechanistically, MAGE-A10 antagonizes the CUL4A-DDB1 E3 ubiquitin ligase complex targeting KAT2A, thereby preventing its K63-linked ubiquitination. Intriguingly, unlike the classic function of MAGEs, which typically assemble with E3 RING ubiquitin ligases to form MAGE-RING ligases (MRLs) and act as a glue for ubiquitination in various cellular processes^37–39^, MAGE-A10 competes with CUL4A-DDB1 for binding to KAT2A at the same interaction region. This binding is also mediated by the MAGE-A10 MHD domain, which typically binds with E3 RING ubiquitin ligases^37,38^. Our results provide a novel regulation mechanism of MAGEs in Cullin-Ring Ubiquitin Ligase (CRL) complexes and expand the understanding of MAGEs’ role in promoting tumorigenesis. Whether other MAGEs share the capacity to bind E3 substrates and prevent substrate degradation will require further investigation.

The mechanisms underlying the aberrant activation of MAGEs in cancers are not yet fully understood. Previous studies suggest that epigenetic dysregulation in cancer cells may trigger their re-expression^61–63^. Although KAT2A does not re-activate MAGE-A10 expression in MAGE-A10-negative cancer cells (data not shown), it significantly boosts MAGE-A10 expression in MAGE-A10-positive cancer cells. The role of KAT2A in enhancing the transcription of MAGE-A10 creates a self-sustaining cycle that perpetuates the aberrant histone acetylation landscape observed in tumors. This feedback loop not only reinforces the stability and activity of KAT2A but also underscores the complexity of epigenetic regulation in cancer. Whether KAT2A also affects the expression of other MAGEs remains to be determined in future studies. Our results indicated that targeting KAT2A inhibition not only impacts its oncogenic function but also fundamentally disrupts the MAGE-A10-KAT2A axis, further eliminating both KAT2A and MAGE-A10 expression.

In summary, our study has unveiled a novel mechanism by which the cancer-testis antigen MAGEA10 orchestrates oncogenic processes. We discovered that MAGEA10 prevents KAT2A from degradation via the autophagy-lysosome pathway by blocking the interaction between KAT2A and the CUL4A-DDB1 E3 ubiquitin ligase complex. The consequent nuclear translocation of KAT2A further amplifies MAGE-A10 transcription, forming a positive feedback loop of MAGE-A10-KAT2A that drives tumor initiation and progression. Therefore, targeting the MAGE-A10-KAT2A axis may offer a novel approach to alter histone acetylation by manipulating KAT2A protein stability for cancer therapy.

## METHODS

### Animals

Male BALB/c nude mice, aged 4-6 weeks and purchased from the GemPharmatech Co., Ltd (Guangdong, China), were used for xenograft growth assays. These mice were maintained in the Laboratory Animal House of the Southern University of Science and Technology, where they were provided with a controlled temperature environment and a 12-hour light/dark cycle. Food and water were available *ad libitum*. All *in vivo* experimental protocols were approved by the Institutional Animal Care and Use Committee (IACUC) of the Southern University of Science and Technology.

### Cell culture

A375, M8, SKMEL-2, HCT116 and HeLa cells were maintained in Dulbecco’s Modified Eagle’s Medium (DMEM) supplemented with 10% FBS. H446, H460, H1975, H1299, H2126 and H1650 cells were maintained in RPMI 1640 medium supplemented with 10% FBS. M059K cells were maintained in DMEM and Ham’s F12 medium (DMEM/F12) supplemented with 10% FBS. U2OS cells were maintained in McCoy’s 5a medium supplemented with 10% FBS. All cells were incubated at 37 °C in a humidified atmosphere containing 5% CO_2_.

### Generation of stable overexpression cell lines

HEK293 cells were transfected with either tandem affinity purification (TAP)-vector or TAP-MAGE-A10 using Effectene in 6 cm^2^ plates. After 48 hours, cells were selected with 1 mg/mL of puromycin over 2 weeks. H460, M8, H1650, H1975, HCT116 and Hela cells were transduced with Myc-vector or Myc-MAGEA10 lentivirus using polybrene in 6-well plates. After stably expressing KAT2A full length (FL) and its mutated (K721R, K728R, K759R) cells in KAT2A/KAT2B-knockout H460 cells, these cells were transduced with Myc-vector or Myc-MAGEA10 lentivirus using polybrene in 6-well plates. Two days after lentiviral transduction, cells were selected over 2 weeks using 2 μg/mL of blasticidin (GIBCO).

### Western blot

Total proteins were separated by SDS-PAGE gel and transferred onto a polyvinylidene fluoride (PVDF) membrane (Millipore, USA) using a Trans-Blot TurboTM (Bio-Rad, USA). Subsequently, the PVDF membrane was blocked with a solution of 5% non-fat dry milk in TBS-T for 1 h at room temperature. After blocking, the membrane was incubated overnight at 4°C with primary antibodies. The membrane was washed three times with TBS-T, followed by incubation with the appropriate HRP-conjugated secondary antibodies (ABclonal) for 1 h at room temperature. The blots were developed using an enhanced chemiluminescence ECL kit (ABclonal) and imaged with a chemiluminescence imaging system (Tanon, Multi5200, CN).

### Antibodies

Anti-MAGE-A10 (Cell Signaling Technology, 68978), anti-GCN5L2 (Cell Signaling Technology, 3305), anti-PCAF (Cell Signaling Technology, 3378), anti-Ubiquitin (Cell Signaling Technology, 20326), anti-Histone H3 (Abcam, ab1791), anti-Histone H3K9 (Abcam, ab4441), anti-Histone H3K14 (Abcam, ab52946), anti-HA tag (Abcam, ab9110), anti-SQSTM1/p62 (Abcam, ab56416), anti-LC3A/B (Abcam, ab62721), anti-CDT2/RAMP (Abcam, ab72264), anti-DDB1 (Abcam, ab109027), anti-LC3 (MBL, M152-3), anti-GCN5L2 (Sigma, SAB1403858), anti-CUL4A (Proteintech, 10693), anti-DDDDK-tag (ABclonal, AE063), anti-MYC-Tag (ABclonal, AE070), anti-His-tag (ABclonal, AE086), anti-c-Myc (ABclonal, A19032), anti-β-Tubulin (ABclonal, A12289), anti-GAPDH (ABclonal, AC002).

### siRNA

siRNA transfections were performed using Lipofectamine RNAiMAX according to the manufacturer’s protocol (Thermo Fisher Scientific). All siRNAs were purchased from OBiO (Shanghai, China) or GenePharma (Suzhou, China). siRNA targeting sequences: siControl, 5’-UUCUCCGAACGUGUCACGUTT, siMAGEA-A10 #1, GCCUGAAGAAGAUCUUCAATT, si-MAGEA-A10 #2, AGUGCAAGUUCUAGCGCUATT, si-MAGEA-A10 #3, GCUGAUGAUGAGACACCAATT; siKAT2A #1 CGAUGUUCGAGCUCUCAAATT, siKAT2A #2, GCUACAUCAAGGACUACGATT siKAT2A #3, CCUCGAAUGAGCAGGUCAATT; siCUL4A, GUUCUUGGACCGCACCUAUTT; siDTL, GCACAUACUUCCAUAGAAATT; sip62, 5’-GAUCUGCGAUGGCUGCAAUTT; siDDB1, ACACGAGAUUAGAGAUCUAUGTT.

### RNA preparation and quantitative reverse transcription PCR Analysis (qRT-PCR)

Total RNA was isolated using the TransZol Up Plus RNA Kit (TranGen). cDNA was synthesized by PrimeScript™ RT reagent Kit (Takara). Gene expression levels were measured by the CFX96 Touch instrument (Bio-Rad, USA) with the Green qPCR SuperMix (TranGen). Gene expression levels were normalized to β-actin and calculated using the 2^−ΔΔCt^ method. Oligonucleotides used for qRT-PCR: MAGEA10: forward: 5’-CAATCCCAAAGTGAGACACAGG, reverse: 5’-GGGGTGCTTGGTATTAGAGGA; KAT2A: forward: 5’-CAGGGTGTGCTGAACTTTGTG, reverse: 5’-TCCAGTAGTTAAGGCAGAGCAA; KAT2B: forward: 5’-AGGAAAACCTGTGGTTGAAGG, reverse: 5’-CAGTCTTCGTTGAGATGGTGC.

### Recombinant protein purification and *in vitro* binding assay

Recombinant protein purification and *in vitro* binding assay were performed as previously described^64^. GST-MAGE-A10, GST-KAT2A, or GST tag alone were expressed in BL21 (DE3) cells at 16°C with isopropyl β-D-1-thiogalactopyranoside (IPTG). Cells were lysed in lysis buffer (50 mM Tris, pH 7.7, 150 mM KCl, 0.1% Triton X-100, 1 mM DTT, 1 mg/mL lysozyme) and the lysates were clarified by centrifugation. GST-tagged proteins were purified using glutathione Sepharose beads (Thermo Fisher Scientific) and eluted with 10 mM glutathione. For *in vitro* binding assay, Myc-tagged proteins were *in vitro* translated using the SP6-TNT Quick rabbit reticulocyte lysate system (Promega) according to the manufacturer’s instructions. The purified GST-tagged proteins bound to glutathione Sepharose beads for 1 hour at 4°C in binding buffer (25 mM Tris, pH 8.0, 2.7 mM KCl, 137 mM NaCl, 0.05% Tween-20, 10 mM 2-mercaptoethanol) and then were blocked for 1 hour in blocking buffer (binding buffer with 5% milk powder). *In vitro* translated Myc-tagged proteins were then added to the beads and incubated for 1 hour at 4°C. After washing, bound proteins were eluted in SDS-sample buffer, boiled, and analyzed by SDS-PAGE, Ponceau S (Beyotime) staining and immunoblotting.

### Tandem ubiquitin binding entity (TUBE) ubiquitination assay

TUBE ubiquitination assay was performed as previously described^65^. A375, H2126, and H460 cells were lysed in TUBE lysis buffer (50 mM Tris, pH 7.5, 150 mM NaCl, 1 mM EDTA, 1% NP-40, 10% glycerol, 20 mM N-Ethylmaleimide (NEM), and 1X protease inhibitor cocktail). The lysates were incubated with magnetic-TUBE beads (LifeSensors) overnight at 4°C. Beads were subsequently washed three times with TBST buffer, and ubiquitinated proteins were eluted in SDS-sample buffer, boiled, and then subjected to SDS-PAGE and immunoblotting.

### Cell viability assay

Cell viability was assessed using a CCK-8 assay (MCE). Cells were seeded in 96-well plates and incubated overnight for adherence. After different treatments, 10 μL of CCK-8 reagent and 90 μL of medium were added to each well. Following a 1.5-hour incubation, absorbance at 450 nm was measured using a PerkinElmer EnSpire® spectrometer.

### Xenograft tumor growth assays

Xenograft tumor growth assays were performed as previously described^66,67^. Wild-type and MAGEA10 knockout cells from A375, SKMEL2 or H446 cells were mixed with matrigel (Corning) before injection of 2 × 10^6^ cells into the flank of BALB/c nude mice. MAGEA10 negative H1975 and M8 cells were made to stably express Myc-vector or Myc-MAGEA10 before injection of 2 × 10^6^ cells in PBS with matrigel into BALB/c nude mice. After stably expressing KAT2A full length (FL) and its mutated (K721R, K728R, K759R) cells in KAT2A/KAT2B-knockout H460 cells, these cells were stably expressed Myc-vector or Myc-MAGEA10 before injection of 2 × 10^6^ cells in PBS with matrigel into BALB/c nude mice. Tumor size was measured every two days during the duration of the experiment. (n = 6 per group).

### CRISPR/Cas9 genome assay

Genetically modified cell lines were generated using CRISPR-Cas9 technology^68^. After cloning guide RNAs (gRNAs) targeting MAGEA10, KAT2A and KAT2B into pCRISPRa_all-in-one. Then these plasmids and packaging plasmids (psPAX2 and pMD2G) were co-transfected into HEK293 cells to generate lentiviral particles using Effectene Transfection Reagent (QIAGEN). A375, SKMEL2, M059K and H446 cells were transfected with lentivirus particles of gRNAs targeting MAGEA10 using polybrene in 6 cm^2^ plates. H460 were transfected with lentivirus particles of gRNAs targeting KAT2A and KAT2B using polybrene in 6 cm^2^ plates. After 48 hours, cells were selected with 4 μg/mL of blasticidin over 1 week. Then, cells were sorted by FACS to enrich for mCherry+ (transfected) cells. The sequences of gRNAs are listed below.

### Table Sequences of gRNAs

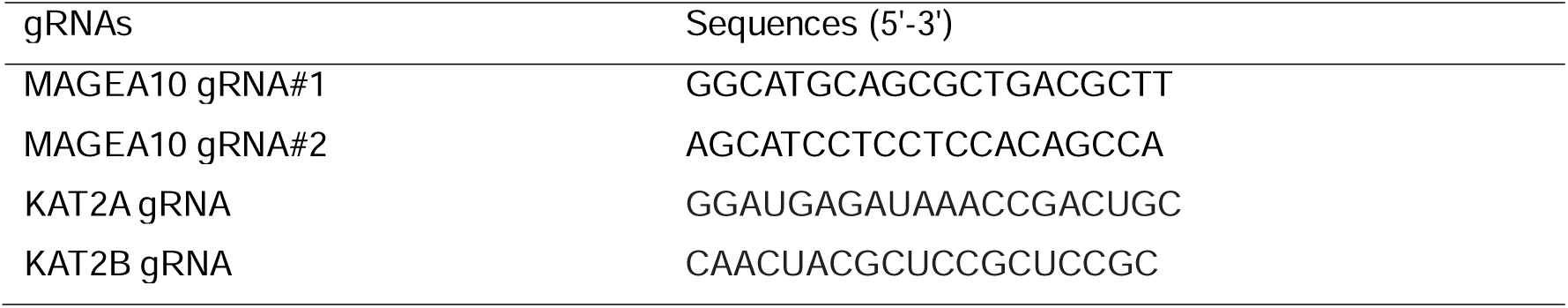

### Colony formation assay

For clonogenic growth assays, wild-type, knockout, or overexpressed cells were seeded in 6-well plates in triplicate. After 1-2 weeks of incubation, the cells were fixed with 10% formalin (Solarbio) and stained with crystal violet (Beyotime) at room temperature for 20 minutes and counted.

### Immunoprecipitation and immunoblotting

Cells were lysed in IP lysis buffer (Thermo Fisher Scientific) supplemented with the protease inhibitor cocktail (NCE). The lysates were centrifuged at 13,000 × g for 10 minutes at 4°C, and the supernatant was incubated with the appropriate primary antibody overnight at 4°C. The antibody-bound proteins were then captured using Protein A/G Magnetic Beads (Thermo Fisher Scientific) for 1-hour incubation at room temperature. To minimize nonspecific binding, the beads were washed thoroughly five times by using a DynaMag™-2 Magnet (Thermo Fisher Scientific, USA). The immunoprecipitated complexes were resuspended in loading buffer, heated at 95°C for 5 minutes, and analyzed by SDS-PAGE followed by western blotting.

### Immunofluorescence staining

Cells were plated onto glass coverslips and fixed with 4% paraformaldehyde in PBS. Following fixation, cells were permeabilized using 0.1% Triton X-100 and blocked with 3% bovine serum albumin (BSA). The cells were then incubated with primary antibodies overnight at 4°C, followed by incubation with secondary antibodies conjugated to (Alexa Fluor 488 or 594 (Abcam). Coverslips are mounted onto microscope slides using an anti-fade mounting medium with DAPI (Abcam). Images were taken with a confocal fluorescence microscope (Zeiss, LSM980, DE).

### *In vitro* ubiquitin assay

To assess the effect of MAGEA10 on KAT2A ubiquitination, *in vitro* ubiquitination assays were conducted following the manufacturer’s protocol (R&D system). The reaction mixture containing ubiquitin (R&D system), MgATP solution (R&D system), E1 enzyme (R&D system), E2 enzyme (R&D system), E3 ligase (CUL4A) (R&D system), E3 ligase reaction buffer (R&D system), KAT2A, and MAGEA10, was incubated at 37°C for 30 minutes. The reaction was terminated by adding 2X protein loading buffer (Epizyme), followed by boiling for 10 minutes. The ubiquitin conjugation reaction products were then analyzed by SDS-PAGE and subsequent western blotting.

### Mass spectrometry analysis

Protein samples were digested and the resulting peptides were analyzed by an optimized LC-MS/MS platform. For quantitative TMT analysis, the digested peptides were labeled with individual TMT reagents, equally pooled, and fractioned by basic pH reversed phase LC chromatography. Each fraction was then analyzed using acidic pH reverse phase nanoscale LC-MS/MS. The collected MS data were processed for protein identification and quantification by database search using the JUMP software suite.

### Statistical analysis

All numerical data are presented as the mean ± SD. A two-sided unpaired t-test was used for comparisons between two groups, while one-way ANOVA followed by Dunnett’s test was applied for comparisons among multiple groups. To assess differences in cell viability and tumor volume between groups, a two-way ANOVA was performed, followed by Tukey’s multiple comparisons tests. *p* < 0.05 was considered statistically significant. All statistical analyses were performed with GraphPad Prism 9 software. We chose the representative images based on the average/median level of the data for each group. For *in vivo* experiments, the sample size was pre-determined by a power calculation. The mice were grouped randomly and blindly by researchers. The mice in poor body condition before the experiments were excluded.

## ACKNOWLEDGMENTS

This work is financially supported by the National Natural Science Foundation of China (No. 82472394 and 82172386 to C.L., and 82100943 to X.F.), the Guangdong Basic and Applied Basic Research Foundation (2022A1515012164 to C.L., and 2023A1515012000 to X.F.), the Science, Technology, and Innovation Commission of Shenzhen (JCYJ20210324104201005 to C.L., and JCYJ20220530115006014 to X.F.), and the Shenzhen Medical Research Fund (A2303061 to X.F.). The authors acknowledge the assistance of the Southern University of Science and Technology Core Research Facilities, the Experimental Animal Center of the Southern University of Science and Technology, and the Clinical Experimental Center, Jiangmen Central Hospital. Fig. 7G was created with BioRender.com.

## AUTHOR CONTRIBUTIONS

X.Y., C.L., and P.R.P. conceptualized the study and designed experiments. X.K.F, J.H and X.Y., performed experiments and analyzed data. X.Y., X.K.F, J.H., and P.R.P. wrote the manuscript.

## DECLARATION OF INTERESTS

The authors declare no competing interests.

## Notes

### Competing Interest Statement

The authors have declared no competing interest.

