## Supplementary material for "A Cancer-Specific Antigen Drives Histone Acetylation by Stabilizing the Acetyltransferases": Figure S1-S7, and Table S1-S2

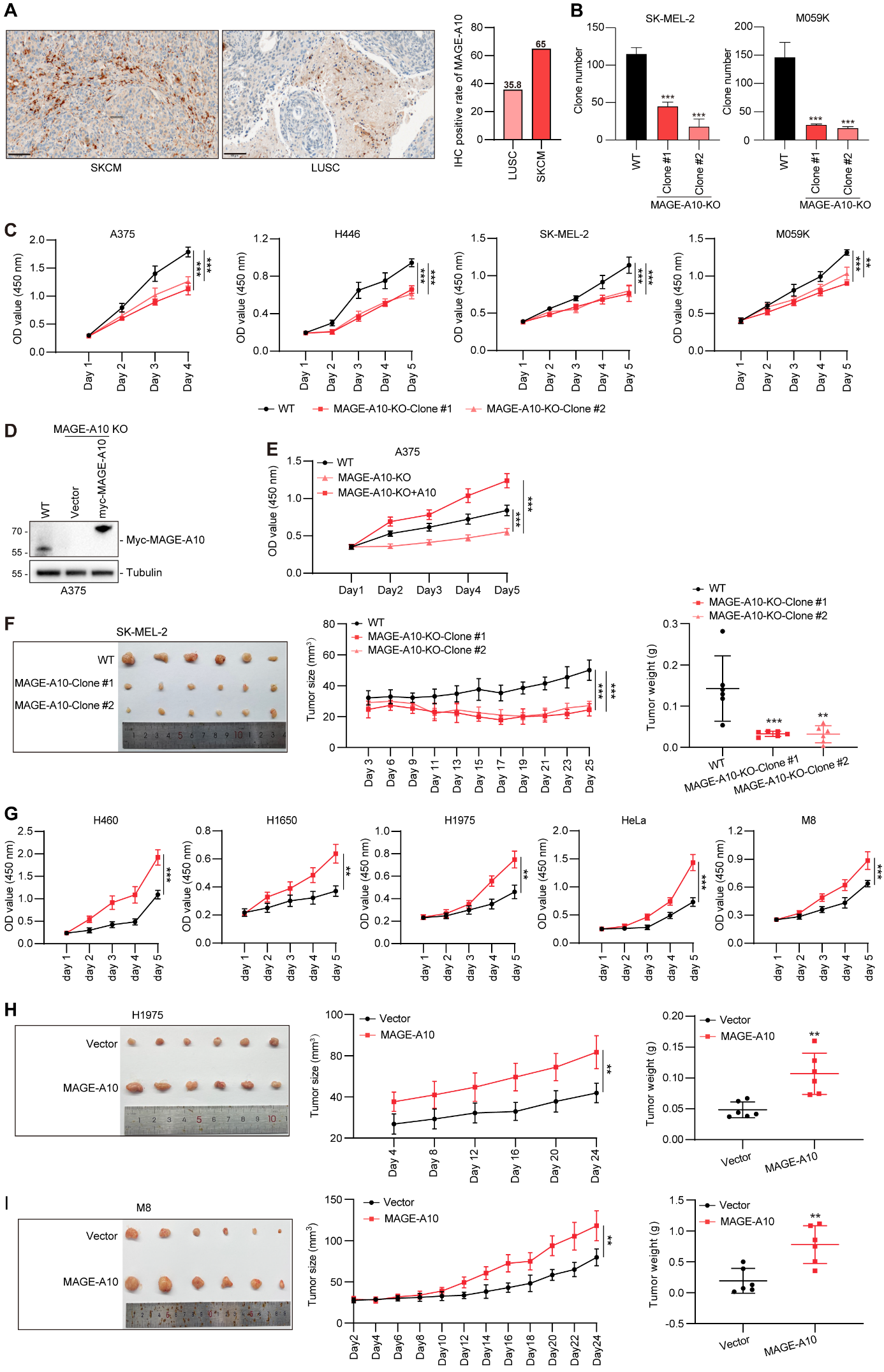

**Figure S1. Aberrant Expression of MAGE-A10 in Cancer Cells Promotes Tumor Growth.**

(A) Immunohistochemical staining of SKCM and LUSC specimens showing MAGE-A10 protein expression in tumor tissues.
(B) Cell viability of wild-type and MAGE-A10-knockout A375, H446, SK-MEL-2, and M059K cells was assessed using the CCK-8 assay.
(C) Clonogenic growth was evaluated in wild-type and MAGE-A10-knockout SK-MEL-2 and M059K clones.
(D) Re-expression of MAGE-A10 in A375 cells was validated through Western blot analysis.
(E) Re-expression of MAGE-A10 restored cell viability in MAGE-A10-knockout A375 cells.
(F) Knockout of MAGE-A10 in SK-MEL-2 cells significantly reduced xenograft tumor growth in mice (n = 6 per group).
(G) The CCK-8 assay was used to evaluate cell viability in H460, H1650, H1975, HeLa and M8 cells with Myc-Vector or Myc-MAGE-A10 re-expression.

(**H-I**) Re-expression of MAGE-A10 in H1975 (H) and M8 (I) cells enhanced xenograft tumor growth in mice (n = 6 per group).

Data are presented as means ± SD. Asterisks denote statistically significant differences compared to controls (*p < 0.05, **p < 0.01, ***p < 0.001, n.s.: not significant).

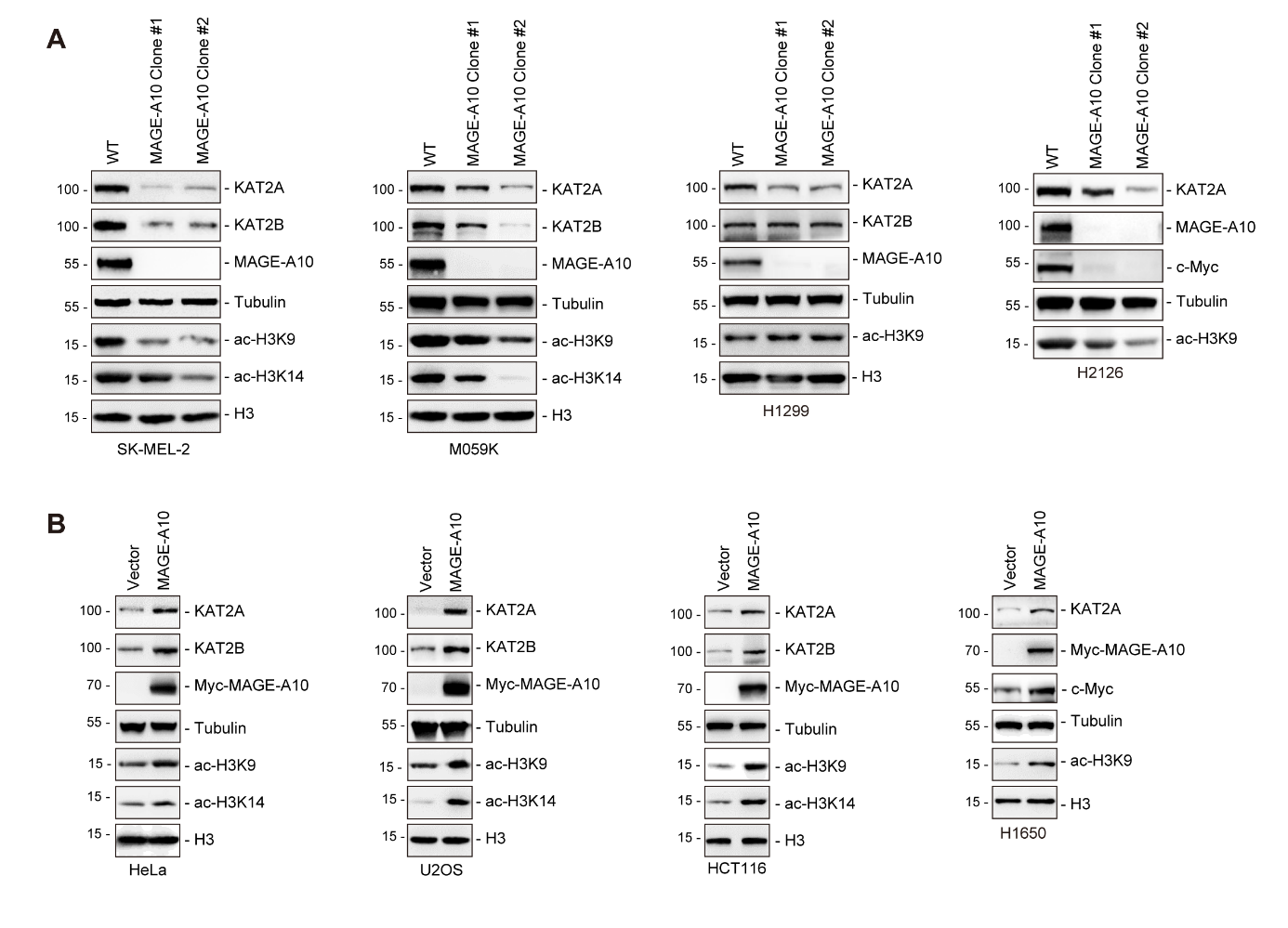

**Figure S2. MAGE-A10 Promotes KAT2A/KAT2B Expression and Histone 3 Lysine Acetylation.**

(A) Knockout of MAGE-A10 decreases KAT2A/KAT2B protein levels and Histone 3 lysine acetylation. Wild-type or MAGE-A10 knockout SK-MEL-2, M059K, H1299 and H2126 cells were blotted for the indicated proteins.
(B) Over-expression of MAGE-A10 increase KAT2A/KAT2B protein levels and Histone 3 lysine acetylation. Indicated proteins were blotted from MYC-vector or MYC-MAGE-A10 over-expressing HeLa, U2OS, HCT116 and H1650 cells, respectively.

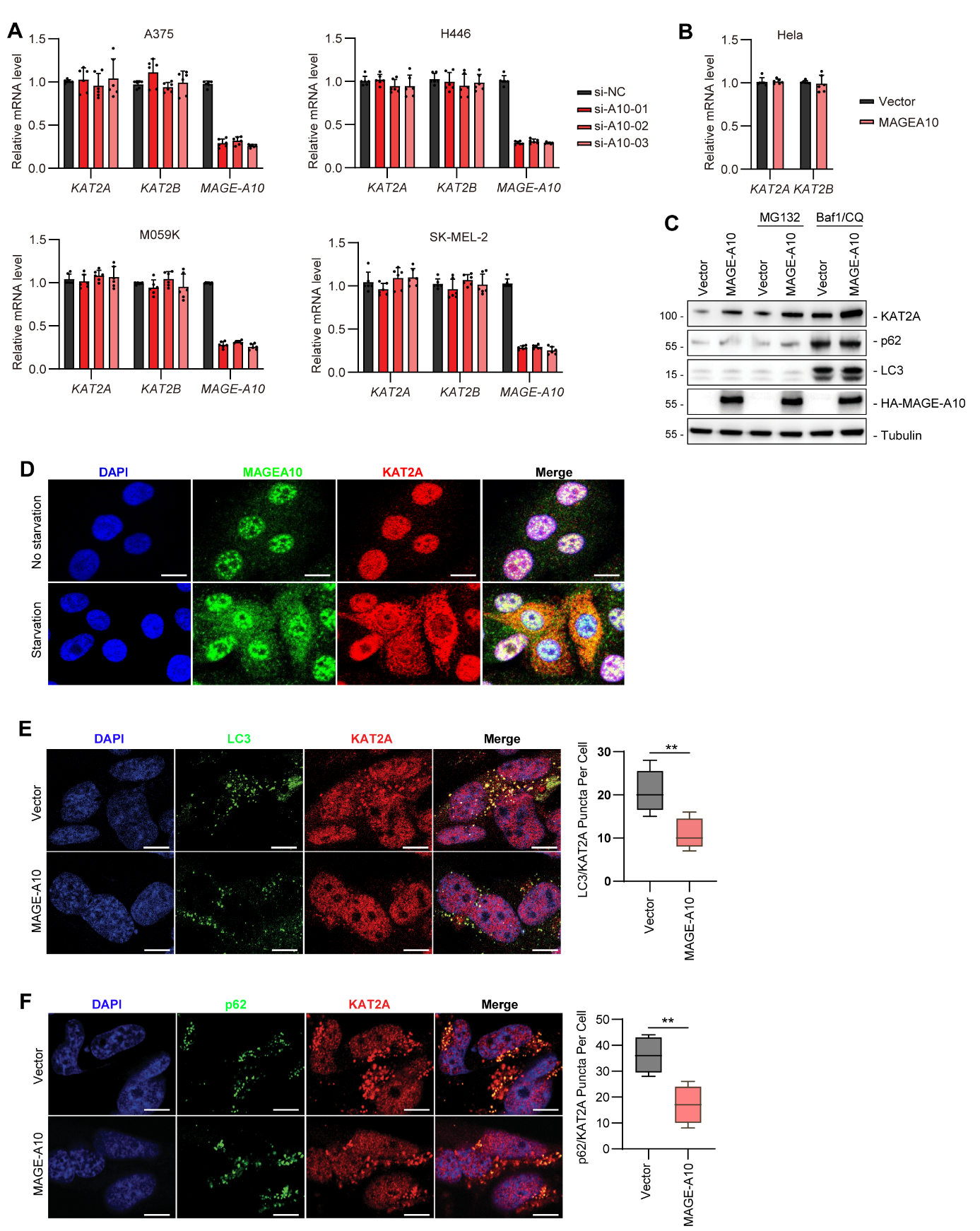

**Figure S3. MAGE-A10 inhibits the autophagic degradation of KAT2A and KAT2B through p62.**

(**A** and **B**) Knockdown or overexpression of MAGE-A10 did not affect KAT2A/KAT2B transcript levels. mRNA levels of KAT2A and KAT2B were measured by qPCR in A375, H446, M059K, and SK-MEL-2 cells treated with si-NC or si-MAGE-A10 (A) and in HeLa cells treated with either a control vector or Myc-MAGE-A10 (B).

(**C**) MAGE-A10 prevented autophagic-dependent KAT2A degradation. A375 cells stably expressing -vector or FLAG-MAGE-A10 were treated with 10 mM MG132 or Baf1 and CQ for 4 h before immunoblotting.

(**D**) Activating autophagy induced the transport of KAT2A into cytoplasm. SK-MEL-2 cells were treated with EBSS with Ca^2+^ and Mg^2+^ prior to immunofluorescence.

(**E**) Over-expression of MAGE-A10 decreased the co-localization of KAT2A with LC3 during autophagy. H460 cells stably expressing MYC-vector or MYC-MAGE-A10 were treated with 200 nM Baf1 and 50 μM CQ for 4 h. Endogenous LC3 and KAT2A were visualized with anti-p62 and anti-KAT2A antibodies staining. The number of KAT2A/LC3 co-localization fluorescent bodies per cell in 40 cells was counted for each data point shown. Scale bar = 10 μm

(**F**) Over-expression of MAGE-A10 decreased the co-localization of KAT2A with p62 during autophagy. H460 cells stably expressing MYC-vector or MYC-MAGE-A10 were treated with 200 nM Baf1 and 50 μM CQ for 4 h. Endogenous p62 and KAT2A were visualized with anti-p62 and anti-KAT2A antibodies staining. The number of KAT2A/p62 co-localization fluorescent bodies per cell in 40 cells was counted for each data point shown. Scale bar = 10 μm

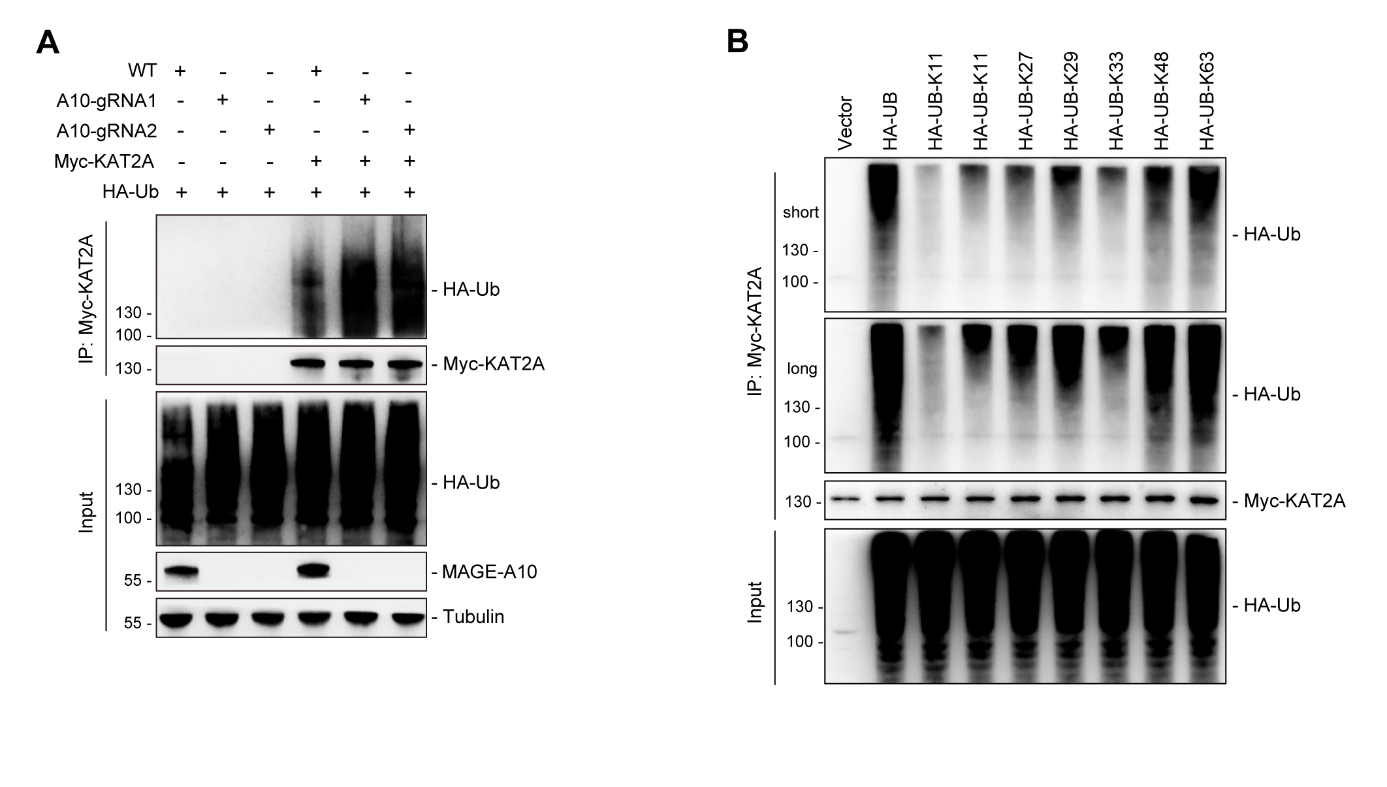

**Figure S4. MAGE-A10 inhibits the K63-linked polyubiquitination of KAT2A by the CUL4A-DDB1 E3 ubiquitin ligase complex.**

(A) Knockout of MAGE-A10 enhanced KAT2A ubiquitination. MAGE-A10 wild-type or knockout M059K cells were transfected with Myc-KAT2A and HA-UB for 72 hours, followed by immunoprecipitation (IP) with anti-Myc-KAT2A antibodies. The indicated proteins were then analyzed by immunoblotting.
(B) Identification of the ubiquitin chain type on KAT2A. HEK293FT cells were transfected with HA-tagged ubiquitin variants (HA-UB, HA-UB-K6, HA-UB-K11, HA-UB-K27, HA-UB-K29, HA-UB-K33, HA-UB-K48, HA-UB-K63) and Myc-KAT2A for 72 hours, followed by immunoprecipitation (IP) with anti-Myc-KAT2A antibodies. The ubiquitin chain types on KAT2A were analyzed by immunoblotting.

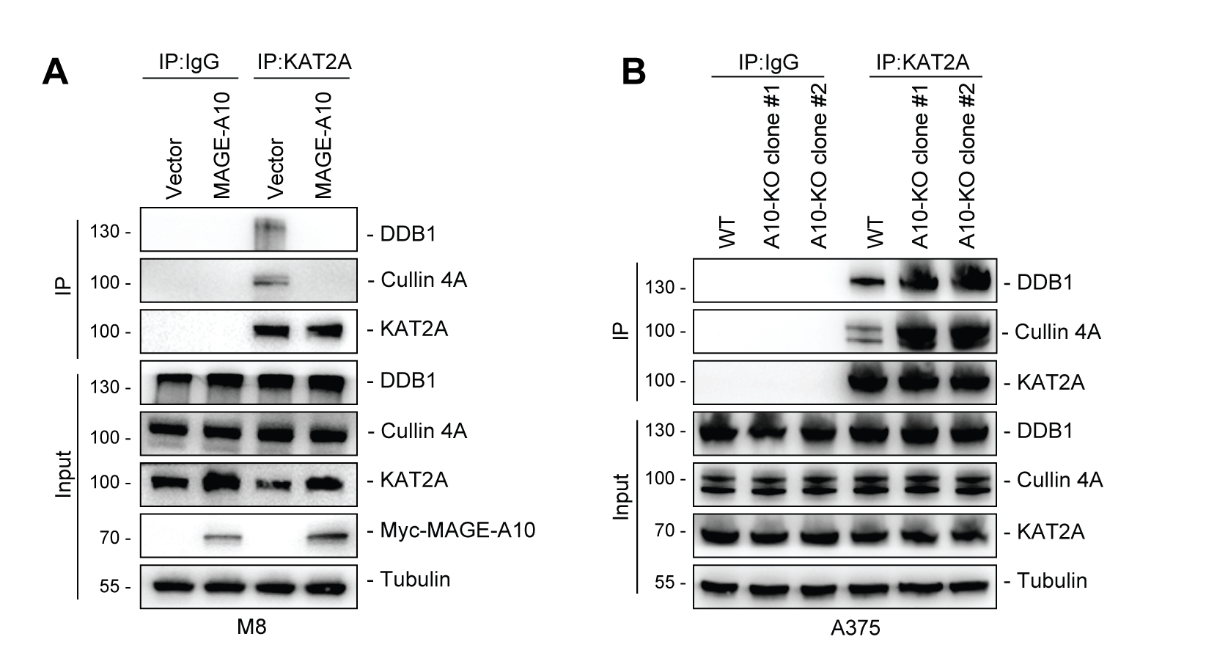

**Figure S5. Cullin 4A-DDB1 mediated KAT2A Ubiquitination was Essential for MAGE-A10-Induced Tumorigenesis.**

(A) MAGE-A10 represses the interaction between KAT2A and the CUL4A-DDB1 complex. Endogenous KAT2A protein was immunoprecipitated from M8 wild-type (WT) or MAGE-A10 overexpressing stable cell lines. The interaction between KAT2A and the CUL4A-DDB1 complex was assessed by immunoblotting.
(B) Deletion of MAGE-A10 increases the interaction between KAT2A and the CUL4A-DDB1 complex. Endogenous KAT2A protein was immunoprecipitated from either A375 wild-type (WT) or with MAGE-A10 knockout stable cell lines. The interaction between KAT2A and the CUL4A-DDB1 complex was examined through immunoblotting.

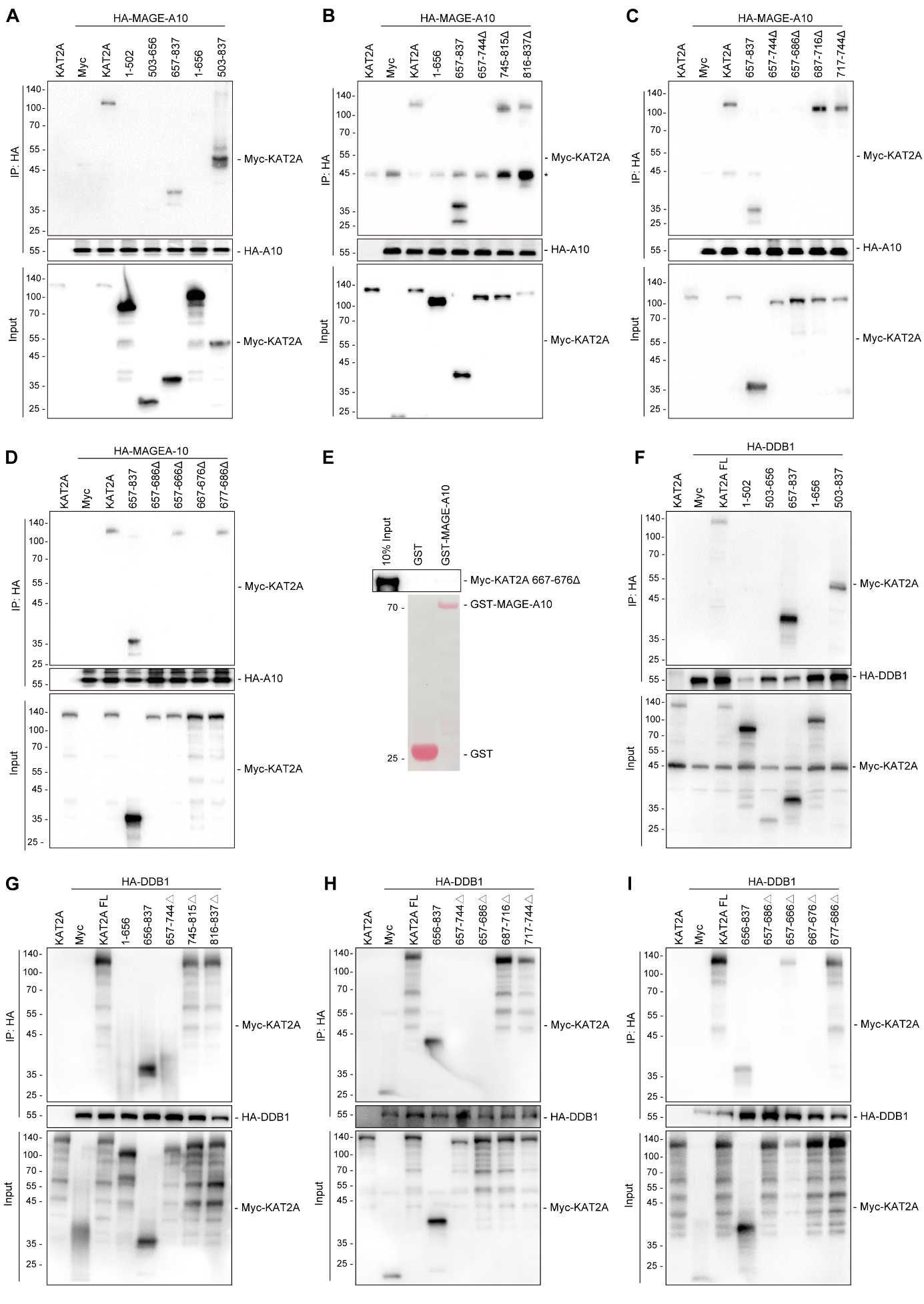

**Figure S6. MAGE-A10 competes with DDB1 for binding to KAT2A.**

(**A-D**) KAT2A has the MAGE-A10 binding region within the sequence 667-676 amino acids. HEK293 cells stably expressing HA-MAGE-A10 were transfected with the indicated constructs for 48 hours before IP with anti-HA followed by SDS-PAGE and immunoblotting for anti-Myc.

(**E-F**) KAT2A 667-676 deletion construct has diminished interaction with MAGE-A10. Myc-tagged KAT2A construct (667-676 deletion) was *in vitro* translated and followed by an *in vitro* binding assay with recombinant GST-MAGE-A10, SDS-PAGE, and immunoblotting for anti-Myc.

(**G-H**) KAT2A has the DDB1 binding region within the sequence 667-676 amino acids. HEK293 cells stably expressing HA-DDB1 were transfected with the indicated constructs for 48 hours before IP with anti-HA followed by SDS-PAGE and immunoblotting for anti-Myc.

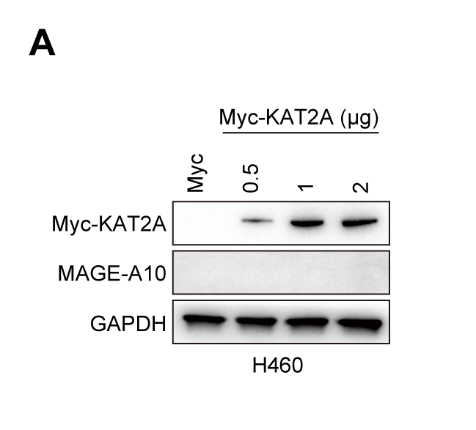

**Figure S7. KAT2A failed to re-activate MAGE-A10 expression in MAGE-A10 negative cancer cells.**

A. Overexpression of KAT2A cannot activate MAGE-A10. H460 cells were transfected with increasing amounts of KAT2A, and MAGE-A10 protein levels were measured by immunoblotting.

**Table S1 MAGE-A10- Spectral count comparison of proteins identified from the Control and MAGE-A10 samples**

| Group | Reference | Description | SC Cntrl2 | SC Magea10 | Mass (KD) | Abundance | p-value |
| --- | --- | --- | --- | --- | --- | --- | --- |
| 115746.1 | sp\|P43363\|MAGAA_HUMAN | Melanoma-associated antigen 10 OS=Homo sapiens GN=MAGEA10 PE=2 SV=2 | 0.1 | 514 | 41 | 1046.48 | 1.3504E-156 |
| 100736.1 | sp\|P08107\|HSP71_HUMAN | Heat shock 70 kDa protein 1A/1B OS=Homo sapiens GN=HSPA1A PE=1 SV=5 | 955 | 192 | 70 | 1691.9 | 1.8632E-122 |
| 115751.1 | sp\|P43358\|MAGA4_HUMAN | Melanoma-associated antigen 4 OS=Homo sapiens GN=MAGEA4 PE=1 SV=2 | 0.1 | 288 | 35 | 954.74 | 1.83555E-88 |
| 115755.1 | sp\|P43362\|MAGA9_HUMAN | Melanoma-associated antigen 9 OS=Homo sapiens GN=MAGEA9 PE=2 SV=1 | 0.1 | 273 | 35 | 1408.73 | 6.14459E-84 |
| 102987.1 | sp\|P34931\|HS71L_HUMAN | Heat shock 70 kDa protein 1-like OS=Homo sapiens GN=HSPA1L PE=1 SV=2 | 366 | 70 | 70 | 713.75 | 8.02853E-50 |
| 110391.1 | sp\|Q9H1R3\|MYLK2_HUMAN | Myosin light chain kinase 2, skeletal/cardiac muscle OS=Homo sapiens GN=MYLK2 PE=1 SV=3 | 8 | 173 | 65 | 230.49 | 3.25313E-42 |
| 103226.1 | sp\|Q86XP3\|DDX42_HUMAN | ATP-dependent RNA helicase DDX42 OS=Homo sapiens GN=DDX42 PE=1 SV=1 | 0.1 | 129 | 103 | 62.67 | 1.84781E-40 |
| 114982.1 | sp\|Q92830\|KAT2A_HUMAN | Histone acetyltransferase KAT2A OS=Homo sapiens GN=KAT2A PE=1 SV=3 | 0.1 | 96 | 94 | 51.14 | 1.78521E-30 |
| 106195.1 | sp\|P39880\|CUX1_HUMAN | Homeobox protein cut-like 1 OS=Homo sapiens GN=CUX1 PE=1 SV=3 | 0.1 | 88 | 164 | 26.81 | 4.73084E-28 |
| 100096.1 | sp\|P08238\|HS90B_HUMAN | Heat shock protein HSP 90-beta OS=Homo sapiens GN=HSP90AB1 PE=1 SV=4 | 2 | 84 | 83 | 72.1 | 1.36142E-23 |
| 107286.1 | sp\|Q15596\|NCOA2_HUMAN | Nuclear receptor coactivator 2 OS=Homo sapiens GN=NCOA2 PE=1 SV=2 | 0.1 | 72 | 159 | 22.63 | 3.35729E-23 |
| 104925.1 | sp\|P49790\|NU153_HUMAN | Nuclear pore complex protein Nup153 OS=Homo sapiens GN=NUP153 PE=1 SV=2 | 0.1 | 51 | 154 | 16.58 | 8.06923E-17 |
| 106195.11 | sp\|Q13948-10\|CASP_HUMAN | Isoform 10 of Protein CASP OS=Homo sapiens GN=CUX1 | 0.1 | 47 | 76 | 31.09 | 1.33332E-15 |
| 100205.1 | sp\|P07900\|HS90A_HUMAN | Heat shock protein HSP 90-alpha OS=Homo sapiens GN=HSP90AA1 PE=1 SV=5 | 0.1 | 44 | 85 | 39.59 | 1.09471E-14 |
| 104965.1 | sp\|P49792\|RBP2_HUMAN | E3 SUMO-protein ligase RanBP2 OS=Homo sapiens GN=RANBP2 PE=1 SV=2 | 0.1 | 44 | 358 | 6.43 | 1.09471E-14 |
| 106195.1 | sp\|Q13948-9\|CASP_HUMAN | Isoform 9 of Protein CASP OS=Homo sapiens GN=CUX1 | 0.1 | 41 | 73 | 28.16 | 9.00418E-14 |
| 101432.1 | sp\|P34932\|HSP74_HUMAN | Heat shock 70 kDa protein 4 OS=Homo sapiens GN=HSPA4 PE=1 SV=4 | 39 | 0.1 | 94 | 33.94 | 3.67309E-13 |
| 114983.1 | sp\|Q92831\|KAT2B_HUMAN | Histone acetyltransferase KAT2B OS=Homo sapiens GN=KAT2B PE=1 SV=3 | 0.1 | 38 | 93 | 20.44 | 7.42142E-13 |
| 107569.1 | sp\|O00541\|PESC_HUMAN | Pescadillo homolog OS=Homo sapiens GN=PES1 PE=1 SV=1 | 0.1 | 36 | 68 | 26.49 | 3.03215E-12 |
| 108636.1 | sp\|O95071\|UBR5_HUMAN | E3 ubiquitin-protein ligase UBR5 OS=Homo sapiens GN=UBR5 PE=1 SV=2 | 0.1 | 33 | 309 | 5.34 | 2.50973E-11 |
| 100112.1 | sp\|P11142\|HSP7C_HUMAN | Heat shock cognate 71 kDa protein OS=Homo sapiens GN=HSPA8 PE=1 SV=1 | 143 | 55 | 71 | 433.28 | 1.95337E-10 |
| 112309.1 | sp\|Q8TD31\|CCHCR_HUMAN | Coiled-coil alpha-helical rod protein 1 OS=Homo sapiens GN=CCHCR1 PE=2 SV=2 | 0.1 | 29 | 89 | 16.36 | 4.22322E-10 |
| 102009.1 | sp\|Q58FF8\|H90B2_HUMAN | Putative heat shock protein HSP 90-beta 2 OS=Homo sapiens GN=HSP90AB2P PE=1 SV=2 | 0.1 | 27 | 44 | 39.48 | 1.73692E-09 |
| 101898.1 | sp\|Q8NBJ4\|GOLM1_HUMAN | Golgi membrane protein 1 OS=Homo sapiens GN=GOLM1 PE=1 SV=1 | 0.1 | 26 | 45 | 28.69 | 3.52513E-09 |
| 102767.6 | sp\|Q9UM54-6\|MYO6_HUMAN | Isoform 6 of Unconventional myosin-VI OS=Homo sapiens GN=MYO6 | 0.1 | 26 | 149 | 8.75 | 3.52513E-09 |
| 106967.1 | sp\|Q9UPW6\|SATB2_HUMAN | DNA-binding protein SATB2 OS=Homo sapiens GN=SATB2 PE=1 SV=2 | 0.1 | 25 | 83 | 15.15 | 7.15836E-09 |
| 113665.1 | sp\|Q8WUB2\|F216A_HUMAN | Protein FAM216A OS=Homo sapiens GN=FAM216A PE=2 SV=1 | 0.1 | 25 | 31 | 40.62 | 7.15836E-09 |
| 101096.1 | sp\|Q92598\|HS105_HUMAN | Heat shock protein 105 kDa OS=Homo sapiens GN=HSPH1 PE=1 SV=1 | 25 | 0.1 | 97 | 19.11 | 7.15836E-09 |
| 102767.3 | sp\|Q9UM54-2\|MYO6_HUMAN | Isoform 2 of Unconventional myosin-VI OS=Homo sapiens GN=MYO6 | 0.1 | 24 | 146 | 8.22 | 1.45449E-08 |
| 107384.1 | sp\|Q9P0K8\|FOXJ2_HUMAN | Forkhead box protein J2 OS=Homo sapiens GN=FOXJ2 PE=1 SV=1 | 0.1 | 24 | 62 | 19.24 | 1.45449E-08 |
| 102767.1 | sp\|Q9UM54\|MYO6_HUMAN | Unconventional myosin-VI OS=Homo sapiens GN=MYO6 PE=1 SV=4 | 0.1 | 23 | 150 | 7.69 | 2.95728E-08 |
| 101195.1 | sp\|P40227\|TCPZ_HUMAN | T-complex protein 1 subunit zeta OS=Homo sapiens GN=CCT6A PE=1 SV=3 | 0.1 | 22 | 58 | 18.97 | 6.01699E-08 |
| 103994.1 | sp\|Q9NXR1\|NDE1_HUMAN | Nuclear distribution protein nudE homolog 1 OS=Homo sapiens GN=NDE1 PE=1 SV=2 | 0.1 | 22 | 39 | 28.36 | 6.01699E-08 |
| 105469.1 | sp\|Q16512\|PKN1_HUMAN | Serine/threonine-protein kinase N1 OS=Homo sapiens GN=PKN1 PE=1 SV=2 | 0.1 | 19 | 104 | 9.15 | 5.09296E-07 |
| 110672.4 | sp\|Q9UPP1-4\|PHF8_HUMAN | Isoform 4 of Histone lysine demethylase PHF8 OS=Homo sapiens GN=PHF8 | 0.1 | 19 | 106 | 8.98 | 5.09296E-07 |
| 100112.1 | tr\|E9PQK7\|E9PQK7_HUMAN | Heat shock cognate 71 kDa protein (Fragment) OS=Homo sapiens GN=HSPA8 PE=1 SV=1 | 59 | 17 | 19 | 603.96 | 7.18234E-07 |
| 102126.1 | sp\|P50991\|TCPD_HUMAN | T-complex protein 1 subunit delta OS=Homo sapiens GN=CCT4 PE=1 SV=4 | 0.1 | 18 | 58 | 15.55 | 1.03991E-06 |
| 110581.1 | sp\|Q99741\|CDC6_HUMAN | Cell division control protein 6 homolog OS=Homo sapiens GN=CDC6 PE=1 SV=1 | 0.1 | 18 | 63 | 14.36 | 1.03991E-06 |
| 104898.1 | sp\|P17066\|HSP76_HUMAN | Heat shock 70 kDa protein 6 OS=Homo sapiens GN=HSPA6 PE=1 SV=2 | 83 | 32 | 71 | 213.42 | 1.2994E-06 |
| 100986.1 | sp\|P62701\|RS4X_HUMAN | 40S ribosomal protein S4, X isoform OS=Homo sapiens GN=RPS4X PE=1 SV=2 | 0.1 | 17 | 30 | 28.74 | 2.12577E-06 |
| 102698.1 | sp\|P46060\|RAGP1_HUMAN | Ran GTPase-activating protein 1 OS=Homo sapiens GN=RANGAP1 PE=1 SV=1 | 0.1 | 17 | 64 | 13.39 | 2.12577E-06 |
| 110672.1 | sp\|Q9UPP1\|PHF8_HUMAN | Histone lysine demethylase PHF8 OS=Homo sapiens GN=PHF8 PE=1 SV=3 | 0.1 | 17 | 118 | 7.22 | 2.12577E-06 |
| 115754.1 | sp\|P43361\|MAGA8_HUMAN | Melanoma-associated antigen 8 OS=Homo sapiens GN=MAGEA8 PE=2 SV=2 | 0.1 | 17 | 35 | 68.19 | 2.12577E-06 |
| 101694.1 | sp\|O14980\|XPO1_HUMAN | Exportin-1 OS=Homo sapiens GN=XPO1 PE=1 SV=1 | 3 | 27 | 123 | 23.92 | 2.60996E-06 |
| 110175.2 | sp\|Q8TD91-2\|MAGC3_HUMAN | Isoform 2 of Melanoma-associated antigen C3 OS=Homo sapiens GN=MAGEC3 | 0.1 | 15 | 39 | 29.58 | 8.91784E-06 |
| 115745.1 | sp\|P43355\|MAGA1_HUMAN | Melanoma-associated antigen 1 OS=Homo sapiens GN=MAGEA1 PE=1 SV=1 | 0.1 | 15 | 34 | 182.1 | 8.91784E-06 |
| 102125.1 | sp\|O95757\|HS74L_HUMAN | Heat shock 70 kDa protein 4L OS=Homo sapiens GN=HSPA4L PE=1 SV=3 | 15 | 0.1 | 94 | 16.94 | 8.91784E-06 |
| 104253.1 | sp\|Q96TA2\|YMEL1_HUMAN | ATP-dependent zinc metalloprotease YME1L1 OS=Homo sapiens GN=YME1L1 PE=1 SV=2 | 0.1 | 14 | 86 | 10.42 | 1.83077E-05 |
| 105738.1 | sp\|P49916\|DNLI3_HUMAN | DNA ligase 3 OS=Homo sapiens GN=LIG3 PE=1 SV=2 | 0.1 | 14 | 113 | 6.2 | 1.83077E-05 |
| 106982.1 | sp\|Q7LGA3\|HS2ST_HUMAN | Heparan sulfate 2-O-sulfotransferase 1 OS=Homo sapiens GN=HS2ST1 PE=1 SV=1 | 0.1 | 14 | 42 | 16.72 | 1.83077E-05 |
| 108939.1 | sp\|Q8IV48\|ERI1_HUMAN | 3'-5' exoribonuclease 1 OS=Homo sapiens GN=ERI1 PE=1 SV=3 | 0.1 | 14 | 40 | 17.48 | 1.83077E-05 |
| 100731.1 | sp\|Q9GZM8\|NDEL1_HUMAN | Nuclear distribution protein nudE-like 1 OS=Homo sapiens GN=NDEL1 PE=1 SV=1 | 0.1 | 13 | 38 | 16.95 | 3.76529E-05 |
| 100220.1 | sp\|P05141\|ADT2_HUMAN | ADP/ATP translocase 2 OS=Homo sapiens GN=SLC25A5 PE=1 SV=7 | 6 | 28 | 33 | 150.77 | 8.48938E-05 |
| 102442.8 | tr\|K7ERQ8\|K7ERQ8_HUMAN | Uncharacterized protein (Fragment) OS=Homo sapiens PE=3 SV=1 | 0.1 | 10 | 31 | 15.94 | 0.000332193 |
| 101718.1 | sp\|Q9UJS0\|CMC2_HUMAN | Calcium-binding mitochondrial carrier protein Aralar2 OS=Homo sapiens GN=SLC25A13 PE=1 SV=2 | 0.1 | 9 | 74 | 30.35 | 0.000690662 |
| 102254.1 | sp\|O60884\|DNJA2_HUMAN | DnaJ homolog subfamily A member 2 OS=Homo sapiens GN=DNAJA2 PE=1 SV=1 | 0.1 | 9 | 46 | 45.93 | 0.000690662 |
| 101513.1 | sp\|P49411\|EFTU_HUMAN | Elongation factor Tu, mitochondrial OS=Homo sapiens GN=TUFM PE=1 SV=2 | 2 | 15 | 50 | 74.73 | 0.000795439 |
| 100190.1 | sp\|P12235\|ADT1_HUMAN | ADP/ATP translocase 1 OS=Homo sapiens GN=SLC25A4 PE=1 SV=4 | 3 | 17 | 33 | 101.38 | 0.001005439 |
| 100223.1 | sp\|P12236\|ADT3_HUMAN | ADP/ATP translocase 3 OS=Homo sapiens GN=SLC25A6 PE=1 SV=4 | 4 | 19 | 33 | 112.65 | 0.001112032 |
| 100976.1 | sp\|Q53H12\|AGK_HUMAN | Acylglycerol kinase, mitochondrial OS=Homo sapiens GN=AGK PE=1 SV=2 | 0.1 | 7 | 47 | 30.78 | 0.003026258 |
| 101306.1 | sp\|Q02978\|M2OM_HUMAN | Mitochondrial 2-oxoglutarate/malate carrier protein OS=Homo sapiens GN=SLC25A11 PE=1 SV=3 | 0.1 | 7 | 34 | 39.66 | 0.003026258 |
| 110942.1 | sp\|P48552\|NRIP1_HUMAN | Nuclear receptor-interacting protein 1 OS=Homo sapiens GN=NRIP1 PE=1 SV=2 | 0.1 | 7 | 127 | 2.76 | 0.003026258 |
| 101157.1 | sp\|Q9NVI7\|ATD3A_HUMAN | ATPase family AAA domain-containing protein 3A OS=Homo sapiens GN=ATAD3A PE=1 SV=2 | 12 | 31 | 71 | 101.65 | 0.003195372 |
| 101157.2 | sp\|Q9NVI7-2\|ATD3A_HUMAN | Isoform 2 of ATPase family AAA domain-containing protein 3A OS=Homo sapiens GN=ATAD3A | 13 | 32 | 66 | 118.62 | 0.004009731 |
| 100935.1 | sp\|Q53GQ0\|DHB12_HUMAN | Estradiol 17-beta-dehydrogenase 12 OS=Homo sapiens GN=HSD17B12 PE=1 SV=2 | 0.1 | 6 | 34 | 21.86 | 0.006393648 |
| 101055.1 | sp\|Q13263\|TIF1B_HUMAN | Transcription intermediary factor 1-beta OS=Homo sapiens GN=TRIM28 PE=1 SV=5 | 0.1 | 6 | 88 | 480.82 | 0.006393648 |
| 101055.2 | sp\|Q13263-2\|TIF1B_HUMAN | Isoform 2 of Transcription intermediary factor 1-beta OS=Homo sapiens GN=TRIM28 | 0.1 | 6 | 79 | 531.32 | 0.006393648 |
| 104910.1 | sp\|Q8NF37\|PCAT1_HUMAN | Lysophosphatidylcholine acyltransferase 1 OS=Homo sapiens GN=LPCAT1 PE=1 SV=2 | 0.1 | 6 | 59 | 18.61 | 0.006393648 |
| 107786.1 | sp\|Q9BQE4\|SELS_HUMAN | Selenoprotein S OS=Homo sapiens GN=VIMP PE=1 SV=3 | 0.1 | 6 | 21 | 14.29 | 0.006393648 |
| 108473.1 | sp\|Q15906\|VPS72_HUMAN | Vacuolar protein sorting-associated protein 72 homolog OS=Homo sapiens GN=VPS72 PE=1 SV=1 | 0.1 | 6 | 41 | 7.39 | 0.006393648 |
| 110925.1 | sp\|Q9BQI6\|ANR32_HUMAN | Ankyrin repeat domain-containing protein 32 OS=Homo sapiens GN=ANKRD32 PE=1 SV=2 | 0.1 | 6 | 121 | 2.48 | 0.006393648 |
| 100138.1 | sp\|Q71U36\|TBA1A_HUMAN | Tubulin alpha-1A chain OS=Homo sapiens GN=TUBA1A PE=1 SV=1 | 6 | 0.1 | 50 | 15.97 | 0.006393648 |
| 100189.1 | sp\|P05023\|AT1A1_HUMAN | Sodium/potassium-transporting ATPase subunit alpha-1 OS=Homo sapiens GN=ATP1A1 PE=1 SV=1 | 0.1 | 5 | 113 | 19.06 | 0.01362796 |
| 100686.1 | sp\|Q9Y230\|RUVB2_HUMAN | RuvB-like 2 OS=Homo sapiens GN=RUVBL2 PE=1 SV=3 | 0.1 | 5 | 51 | 16.63 | 0.01362796 |
| 101799.1 | sp\|Q9NZ01\|TECR_HUMAN | Very-long-chain enoyl-CoA reductase OS=Homo sapiens GN=TECR PE=1 SV=1 | 0.1 | 5 | 36 | 27.77 | 0.01362796 |
| 103842.1 | sp\|Q9NPF5\|DMAP1_HUMAN | DNA methyltransferase 1-associated protein 1 OS=Homo sapiens GN=DMAP1 PE=1 SV=1 | 0.1 | 5 | 53 | 4.72 | 0.01362796 |
| 104585.1 | sp\|P29590\|PML_HUMAN | Protein PML OS=Homo sapiens GN=PML PE=1 SV=3 | 0.1 | 5 | 97 | 2.56 | 0.01362796 |
| 105705.1 | sp\|P43307\|SSRA_HUMAN | Translocon-associated protein subunit alpha OS=Homo sapiens GN=SSR1 PE=1 SV=3 | 0.1 | 5 | 32 | 32.59 | 0.01362796 |
| 109316.3 | sp\|Q9ULL5-3\|PRR12_HUMAN | Isoform 3 of Proline-rich protein 12 OS=Homo sapiens GN=PRR12 | 0.1 | 5 | 211 | 1.19 | 0.01362796 |
| 105894.1 | sp\|Q07021\|C1QBP_HUMAN | Complement component 1 Q subcomponent-binding protein, mitochondrial OS=Homo sapiens GN=C1QBP PE=1 SV=1 | 5 | 0.1 | 31 | 28.71 | 0.01362796 |
| 100331.1 | sp\|Q9BVA1\|TBB2B_HUMAN | Tubulin beta-2B chain OS=Homo sapiens GN=TUBB2B PE=1 SV=1 | 29 | 50 | 50 | 331.52 | 0.017455845 |
| 100094.1 | sp\|Q13885\|TBB2A_HUMAN | Tubulin beta-2A chain OS=Homo sapiens GN=TUBB2A PE=1 SV=1 | 30 | 51 | 50 | 337.84 | 0.018939017 |
| 119534.1 | sp\|P04350\|TBB4A_HUMAN | Tubulin beta-4A chain OS=Homo sapiens GN=TUBB4A PE=1 SV=2 | 31 | 52 | 50 | 377.36 | 0.02046902 |
| 119535.1 | sp\|P68371\|TBB4B_HUMAN | Tubulin beta-4B chain OS=Homo sapiens GN=TUBB4B PE=1 SV=1 | 41 | 64 | 50 | 444.78 | 0.02421484 |
| 100563.1 | sp\|O75746\|CMC1_HUMAN | Calcium-binding mitochondrial carrier protein Aralar1 OS=Homo sapiens GN=SLC25A12 PE=1 SV=2 | 0.1 | 4 | 75 | 4.02 | 0.0294083 |
| 100853.1 | sp\|P14625\|ENPL_HUMAN | Endoplasmin OS=Homo sapiens GN=HSP90B1 PE=1 SV=1 | 0.1 | 4 | 92 | 7.57 | 0.0294083 |
| 100864.1 | sp\|O95831\|AIFM1_HUMAN | Apoptosis-inducing factor 1, mitochondrial OS=Homo sapiens GN=AIFM1 PE=1 SV=1 | 0.1 | 4 | 67 | 20.94 | 0.0294083 |
| 101062.1 | sp\|Q9BXF6\|RFIP5_HUMAN | Rab11 family-interacting protein 5 OS=Homo sapiens GN=RAB11FIP5 PE=1 SV=1 | 0.1 | 4 | 70 | 2.84 | 0.0294083 |
| 101384.1 | sp\|P02794\|FRIH_HUMAN | Ferritin heavy chain OS=Homo sapiens GN=FTH1 PE=1 SV=2 | 0.1 | 4 | 21 | 42.43 | 0.0294083 |
| 101408.1 | sp\|P33176\|KINH_HUMAN | Kinesin-1 heavy chain OS=Homo sapiens GN=KIF5B PE=1 SV=1 | 0.1 | 4 | 110 | 1.82 | 0.0294083 |
| 101483.1 | sp\|P51648\|AL3A2_HUMAN | Fatty aldehyde dehydrogenase OS=Homo sapiens GN=ALDH3A2 PE=1 SV=1 | 0.1 | 4 | 55 | 3.65 | 0.0294083 |
| 101694.1 | tr\|C9JF49\|C9JF49_HUMAN | Exportin-1 (Fragment) OS=Homo sapiens GN=XPO1 PE=1 SV=1 | 0.1 | 4 | 7 | 67.64 | 0.0294083 |
| 102107.1 | sp\|Q8N2K0\|ABD12_HUMAN | Monoacylglycerol lipase ABHD12 OS=Homo sapiens GN=ABHD12 PE=1 SV=2 | 0.1 | 4 | 45 | 7.77 | 0.0294083 |
| 103556.1 | sp\|Q15029\|U5S1_HUMAN | 116 kDa U5 small nuclear ribonucleoprotein component OS=Homo sapiens GN=EFTUD2 PE=1 SV=1 | 0.1 | 4 | 109 | 10.97 | 0.0294083 |
| 104585.1 | sp\|P29590-12\|PML_HUMAN | Isoform PML-12 of Protein PML OS=Homo sapiens GN=PML | 0.1 | 4 | 65 | 3.08 | 0.0294083 |
| 105453.1 | sp\|Q96N67\|DOCK7_HUMAN | Dedicator of cytokinesis protein 7 OS=Homo sapiens GN=DOCK7 PE=1 SV=4 | 0.1 | 4 | 242 | 0.83 | 0.0294083 |
| 109316.1 | sp\|Q9ULL5\|PRR12_HUMAN | Proline-rich protein 12 OS=Homo sapiens GN=PRR12 PE=1 SV=2 | 0.1 | 4 | 130 | 1.54 | 0.0294083 |
| 115838.1 | sp\|P33993\|MCM7_HUMAN | DNA replication licensing factor MCM7 OS=Homo sapiens GN=MCM7 PE=1 SV=4 | 0.1 | 4 | 81 | 14.77 | 0.0294083 |
| 100283.1 | sp\|P55072\|TERA_HUMAN | Transitional endoplasmic reticulum ATPase OS=Homo sapiens GN=VCP PE=1 SV=4 | 4 | 0.1 | 89 | 17.36 | 0.0294083 |
| 101944.1 | sp\|P0CG05\|LAC2_HUMAN | Ig lambda-2 chain C regions OS=Homo sapiens GN=IGLC2 PE=1 SV=1 | 4 | 0.1 | 11 | 101.88 | 0.0294083 |
| 102746.1 | sp\|Q15365\|PCBP1_HUMAN | Poly(rC)-binding protein 1 OS=Homo sapiens GN=PCBP1 PE=1 SV=2 | 4 | 0.1 | 37 | 5.34 | 0.0294083 |
| 103593.1 | sp\|Q9NPH2\|INO1_HUMAN | Inositol-3-phosphate synthase 1 OS=Homo sapiens GN=ISYNA1 PE=1 SV=1 | 4 | 0.1 | 61 | 16.39 | 0.0294083 |
| 105571.1 | sp\|Q9NTJ4\|MA2C1_HUMAN | Alpha-mannosidase 2C1 OS=Homo sapiens GN=MAN2C1 PE=1 SV=1 | 4 | 0.1 | 116 | 10.37 | 0.0294083 |
| 100108.1 | sp\|P07437\|TBB5_HUMAN | Tubulin beta chain OS=Homo sapiens GN=TUBB PE=1 SV=2 | 46 | 69 | 50 | 467.37 | 0.031396181 |
| 100413.1 | sp\|P11021\|GRP78_HUMAN | 78 kDa glucose-regulated protein OS=Homo sapiens GN=HSPA5 PE=1 SV=2 | 24 | 41 | 72 | 131.42 | 0.033928931 |
| 100738.1 | sp\|P54652\|HSP72_HUMAN | Heat shock-related 70 kDa protein 2 OS=Homo sapiens GN=HSPA2 PE=1 SV=1 | 37 | 21 | 70 | 132.9 | 0.034463681 |
| 102682.1 | sp\|Q5T9A4\|ATD3B_HUMAN | ATPase family AAA domain-containing protein 3B OS=Homo sapiens GN=ATAD3B PE=1 SV=1 | 11 | 22 | 73 | 73.07 | 0.053195115 |
| 102741.1 | sp\|Q3ZCM7\|TBB8_HUMAN | Tubulin beta-8 chain OS=Homo sapiens GN=TUBB8 PE=1 SV=2 | 11 | 22 | 50 | 152.78 | 0.053195115 |
| 100371.1 | sp\|Q01813\|PFKAP_HUMAN | ATP-dependent 6-phosphofructokinase, platelet type OS=Homo sapiens GN=PFKP PE=1 SV=2 | 0.1 | 3 | 86 | 13.44 | 0.064646504 |
| 100662.1 | sp\|P27361\|MK03_HUMAN | Mitogen-activated protein kinase 3 OS=Homo sapiens GN=MAPK3 PE=1 SV=4 | 0.1 | 3 | 43 | 12.76 | 0.064646504 |
| 100723.1 | sp\|Q9H936\|GHC1_HUMAN | Mitochondrial glutamate carrier 1 OS=Homo sapiens GN=SLC25A22 PE=1 SV=1 | 0.1 | 3 | 34 | 14.51 | 0.064646504 |
| 101572.1 | sp\|P16615\|AT2A2_HUMAN | Sarcoplasmic/endoplasmic reticulum calcium ATPase 2 OS=Homo sapiens GN=ATP2A2 PE=1 SV=1 | 0.1 | 3 | 115 | 9.16 | 0.064646504 |
| 101798.1 | sp\|P36542\|ATPG_HUMAN | ATP synthase subunit gamma, mitochondrial OS=Homo sapiens GN=ATP5C1 PE=1 SV=1 | 0.1 | 3 | 33 | 24.26 | 0.064646504 |
| 102149.1 | sp\|P53985\|MOT1_HUMAN | Monocarboxylate transporter 1 OS=Homo sapiens GN=SLC16A1 PE=1 SV=3 | 0.1 | 3 | 54 | 8.35 | 0.064646504 |
| 102273.1 | sp\|Q9Y2X3\|NOP58_HUMAN | Nucleolar protein 58 OS=Homo sapiens GN=NOP58 PE=1 SV=1 | 0.1 | 3 | 60 | 20.15 | 0.064646504 |
| 102838.1 | sp\|Q8TAQ2\|SMRC2_HUMAN | SWI/SNF complex subunit SMARCC2 OS=Homo sapiens GN=SMARCC2 PE=1 SV=1 | 0.1 | 3 | 133 | 1.13 | 0.064646504 |
| 102886.1 | sp\|Q96QK1\|VPS35_HUMAN | Vacuolar protein sorting-associated protein 35 OS=Homo sapiens GN=VPS35 PE=1 SV=2 | 0.1 | 3 | 92 | 10.91 | 0.064646504 |
| 103504.1 | sp\|P61619\|S61A1_HUMAN | Protein transport protein Sec61 subunit alpha isoform 1 OS=Homo sapiens GN=SEC61A1 PE=1 SV=2 | 0.1 | 3 | 52 | 11.49 | 0.064646504 |
| 103802.1 | sp\|Q9Y5M8\|SRPRB_HUMAN | Signal recognition particle receptor subunit beta OS=Homo sapiens GN=SRPRB PE=1 SV=3 | 0.1 | 3 | 30 | 8.42 | 0.064646504 |
| 104042.1 | sp\|P63165\|SUMO1_HUMAN | Small ubiquitin-related modifier 1 OS=Homo sapiens GN=SUMO1 PE=1 SV=1 | 0.1 | 3 | 12 | 12.99 | 0.064646504 |
| 104354.1 | sp\|O96019\|ACL6A_HUMAN | Actin-like protein 6A OS=Homo sapiens GN=ACTL6A PE=1 SV=1 | 0.1 | 3 | 47 | 3.16 | 0.064646504 |
| 106199.1 | sp\|Q6UUV7\|CRTC3_HUMAN | CREB-regulated transcription coactivator 3 OS=Homo sapiens GN=CRTC3 PE=1 SV=2 | 0.1 | 3 | 67 | 2.24 | 0.064646504 |
| 106431.1 | sp\|Q9Y618\|NCOR2_HUMAN | Nuclear receptor corepressor 2 OS=Homo sapiens GN=NCOR2 PE=1 SV=2 | 0.1 | 3 | 275 | 0.55 | 0.064646504 |
| 106673.1 | sp\|Q9NVM6\|DJC17_HUMAN | DnaJ homolog subfamily C member 17 OS=Homo sapiens GN=DNAJC17 PE=1 SV=1 | 0.1 | 3 | 35 | 4.33 | 0.064646504 |
| 106984.1 | sp\|Q6R327\|RICTR_HUMAN | Rapamycin-insensitive companion of mTOR OS=Homo sapiens GN=RICTOR PE=1 SV=1 | 0.1 | 3 | 192 | 0.78 | 0.064646504 |
| 107456.1 | sp\|Q9Y2J4\|AMOL2_HUMAN | Angiomotin-like protein 2 OS=Homo sapiens GN=AMOTL2 PE=1 SV=3 | 0.1 | 3 | 86 | 1.75 | 0.064646504 |
| 108824.1 | sp\|Q6ZRS2\|SRCAP_HUMAN | Helicase SRCAP OS=Homo sapiens GN=SRCAP PE=1 SV=3 | 0.1 | 3 | 343 | 0.44 | 0.064646504 |
| 115553.1 | sp\|Q8N9Z9\|LMTD1_HUMAN | Lamin tail domain-containing protein 1 OS=Homo sapiens GN=LMNTD1 PE=2 SV=2 | 0.1 | 3 | 43 | 3.46 | 0.064646504 |
| 118176.1 | sp\|Q15758\|AAAT_HUMAN | Neutral amino acid transporter B(0) OS=Homo sapiens GN=SLC1A5 PE=1 SV=2 | 0.1 | 3 | 57 | 2.65 | 0.064646504 |
| 102402.1 | tr\|A0A087WXC3\|A0A087WXC3_HUMAN | Immunoglobulin lambda-like polypeptide 5 OS=Homo sapiens GN=IGLL5 PE=4 SV=1 | 3 | 0.1 | 25 | 30.33 | 0.064646504 |
| 107301.1 | sp\|P52701\|MSH6_HUMAN | DNA mismatch repair protein Msh6 OS=Homo sapiens GN=MSH6 PE=1 SV=2 | 3 | 0.1 | 153 | 4.26 | 0.064646504 |
| 101099.2 | sp\|Q00325-2\|MPCP_HUMAN | Isoform B of Phosphate carrier protein, mitochondrial OS=Homo sapiens GN=SLC25A3 | 3 | 9 | 40 | 68.86 | 0.076417753 |
| 100202.1 | sp\|P23396\|RS3_HUMAN | 40S ribosomal protein S3 OS=Homo sapiens GN=RPS3 PE=1 SV=2 | 5 | 12 | 27 | 136.85 | 0.084827666 |
| 101099.1 | sp\|Q00325\|MPCP_HUMAN | Phosphate carrier protein, mitochondrial OS=Homo sapiens GN=SLC25A3 PE=1 SV=2 | 2 | 7 | 40 | 53.66 | 0.086307142 |
| 100217.1 | sp\|P25705\|ATPA_HUMAN | ATP synthase subunit alpha, mitochondrial OS=Homo sapiens GN=ATP5A1 PE=1 SV=1 | 11 | 20 | 60 | 108.85 | 0.103490933 |
| 102165.1 | sp\|P01857\|IGHG1_HUMAN | Ig gamma-1 chain C region OS=Homo sapiens GN=IGHG1 PE=1 SV=1 | 15 | 8 | 36 | 199.53 | 0.141220717 |
| 102165.4 | tr\|A0A087X079\|A0A087X079_HUMAN | Ig gamma-1 chain C region OS=Homo sapiens GN=IGHG1 PE=1 SV=1 | 15 | 8 | 52 | 139.74 | 0.141220717 |
| 100058.1 | sp\|P42677\|RS27_HUMAN | 40S ribosomal protein S27 OS=Homo sapiens GN=RPS27 PE=1 SV=3 | 0.1 | 2 | 9 | 68.74 | 0.146611857 |
| 100146.1 | sp\|P62269\|RS18_HUMAN | 40S ribosomal protein S18 OS=Homo sapiens GN=RPS18 PE=1 SV=3 | 0.1 | 2 | 18 | 22.59 | 0.146611857 |
| 100168.1 | sp\|P13637\|AT1A3_HUMAN | Sodium/potassium-transporting ATPase subunit alpha-3 OS=Homo sapiens GN=ATP1A3 PE=1 SV=3 | 0.1 | 2 | 112 | 7.16 | 0.146611857 |
| 100208.1 | sp\|P10809\|CH60_HUMAN | 60 kDa heat shock protein, mitochondrial OS=Homo sapiens GN=HSPD1 PE=1 SV=2 | 0.1 | 2 | 61 | 8.19 | 0.146611857 |
| 100487.1 | sp\|P06576\|ATPB_HUMAN | ATP synthase subunit beta, mitochondrial OS=Homo sapiens GN=ATP5B PE=1 SV=3 | 0.1 | 2 | 57 | 12.38 | 0.146611857 |
| 100681.11 | tr\|E9PF82\|E9PF82_HUMAN | Calcium/calmodulin-dependent protein kinase type II subunit delta OS=Homo sapiens GN=CAMK2D PE=1 SV=1 | 0.1 | 2 | 60 | 1.67 | 0.146611857 |
| 100688.1 | sp\|P49368\|TCPG_HUMAN | T-complex protein 1 subunit gamma OS=Homo sapiens GN=CCT3 PE=1 SV=4 | 0.1 | 2 | 60 | 6.61 | 0.146611857 |
| 100721.1 | sp\|O60282\|KIF5C_HUMAN | Kinesin heavy chain isoform 5C OS=Homo sapiens GN=KIF5C PE=1 SV=1 | 0.1 | 2 | 109 | 0.91 | 0.146611857 |
| 100771.1 | sp\|Q9ULA0\|DNPEP_HUMAN | Aspartyl aminopeptidase OS=Homo sapiens GN=DNPEP PE=1 SV=1 | 0.1 | 2 | 52 | 3.82 | 0.146611857 |
| 100861.3 | tr\|E9PCA1\|E9PCA1_HUMAN | T-complex protein 1 subunit epsilon OS=Homo sapiens GN=CCT5 PE=1 SV=1 | 0.1 | 2 | 57 | 1.75 | 0.146611857 |
| 101395.1 | sp\|O75122\|CLAP2_HUMAN | CLIP-associating protein 2 OS=Homo sapiens GN=CLASP2 PE=1 SV=2 | 0.1 | 2 | 141 | 0.71 | 0.146611857 |
| 101461.1 | sp\|P35613\|BASI_HUMAN | Basigin OS=Homo sapiens GN=BSG PE=1 SV=2 | 0.1 | 2 | 42 | 10.67 | 0.146611857 |
| 101503.1 | sp\|P26641\|EF1G_HUMAN | Elongation factor 1-gamma OS=Homo sapiens GN=EEF1G PE=1 SV=3 | 0.1 | 2 | 50 | 24.96 | 0.146611857 |
| 101833.1 | sp\|O43175\|SERA_HUMAN | D-3-phosphoglycerate dehydrogenase OS=Homo sapiens GN=PHGDH PE=1 SV=4 | 0.1 | 2 | 57 | 15.01 | 0.146611857 |
| 102081.1 | sp\|P08579\|RU2B_HUMAN | U2 small nuclear ribonucleoprotein B'' OS=Homo sapiens GN=SNRPB2 PE=1 SV=1 | 0.1 | 2 | 25 | 3.93 | 0.146611857 |
| 102089.1 | sp\|P50454\|SERPH_HUMAN | Serpin H1 OS=Homo sapiens GN=SERPINH1 PE=1 SV=2 | 0.1 | 2 | 46 | 15.08 | 0.146611857 |
| 102196.1 | sp\|P08574\|CY1_HUMAN | Cytochrome c1, heme protein, mitochondrial OS=Homo sapiens GN=CYC1 PE=1 SV=3 | 0.1 | 2 | 35 | 2.82 | 0.146611857 |
| 102644.1 | sp\|Q9UPW0\|FOXJ3_HUMAN | Forkhead box protein J3 OS=Homo sapiens GN=FOXJ3 PE=1 SV=2 | 0.1 | 2 | 69 | 1.45 | 0.146611857 |
| 102726.1 | tr\|E7ENN3\|E7ENN3_HUMAN | Nesprin-1 OS=Homo sapiens GN=SYNE1 PE=1 SV=2 | 0.1 | 2 | 964 | 0.1 | 0.146611857 |
| 102918.1 | sp\|Q969G3\|SMCE1_HUMAN | SWI/SNF-related matrix-associated actin-dependent regulator of chromatin subfamily E member 1 OS=Homo sapiens GN=SMARCE1 PE=1 SV=2 | 0.1 | 2 | 47 | 2.14 | 0.146611857 |
| 103109.1 | sp\|Q92499\|DDX1_HUMAN | ATP-dependent RNA helicase DDX1 OS=Homo sapiens GN=DDX1 PE=1 SV=2 | 0.1 | 2 | 82 | 8.5 | 0.146611857 |
| 103123.1 | sp\|O00159\|MYO1C_HUMAN | Unconventional myosin-Ic OS=Homo sapiens GN=MYO1C PE=1 SV=4 | 0.1 | 2 | 122 | 0.82 | 0.146611857 |
| 103994.1 | tr\|I3L464\|I3L464_HUMAN | Nuclear distribution protein nudE homolog 1 OS=Homo sapiens GN=NDE1 PE=4 SV=1 | 0.1 | 2 | 13 | 7.7 | 0.146611857 |
| 105939.1 | sp\|Q7L5N7\|PCAT2_HUMAN | Lysophosphatidylcholine acyltransferase 2 OS=Homo sapiens GN=LPCAT2 PE=1 SV=1 | 0.1 | 2 | 60 | 1.66 | 0.146611857 |
| 106516.1 | sp\|Q92621\|NU205_HUMAN | Nuclear pore complex protein Nup205 OS=Homo sapiens GN=NUP205 PE=1 SV=3 | 0.1 | 2 | 228 | 4.17 | 0.146611857 |
| 106570.1 | sp\|Q5H9J7\|BEX5_HUMAN | Protein BEX5 OS=Homo sapiens GN=BEX5 PE=2 SV=1 | 0.1 | 2 | 13 | 7.94 | 0.146611857 |
| 106749.1 | sp\|O75151\|PHF2_HUMAN | Lysine-specific demethylase PHF2 OS=Homo sapiens GN=PHF2 PE=1 SV=4 | 0.1 | 2 | 121 | 0.83 | 0.146611857 |
| 106781.1 | sp\|Q9HD23\|MRS2_HUMAN | Magnesium transporter MRS2 homolog, mitochondrial OS=Homo sapiens GN=MRS2 PE=1 SV=1 | 0.1 | 2 | 50 | 1.99 | 0.146611857 |
| 106793.1 | sp\|Q8IVF7\|FMNL3_HUMAN | Formin-like protein 3 OS=Homo sapiens GN=FMNL3 PE=1 SV=3 | 0.1 | 2 | 117 | 0.85 | 0.146611857 |
| 106945.1 | sp\|Q9BWH6\|RPAP1_HUMAN | RNA polymerase II-associated protein 1 OS=Homo sapiens GN=RPAP1 PE=1 SV=3 | 0.1 | 2 | 153 | 0.66 | 0.146611857 |
| 108368.1 | sp\|Q92922\|SMRC1_HUMAN | SWI/SNF complex subunit SMARCC1 OS=Homo sapiens GN=SMARCC1 PE=1 SV=3 | 0.1 | 2 | 123 | 0.81 | 0.146611857 |
| 119411.1 | sp\|O15164\|TIF1A_HUMAN | Transcription intermediary factor 1-alpha OS=Homo sapiens GN=TRIM24 PE=1 SV=3 | 0.1 | 2 | 117 | 0.86 | 0.146611857 |
| 100025.2 | tr\|M0QZ52\|M0QZ52_HUMAN | Calmodulin OS=Homo sapiens GN=CALM3 PE=1 SV=1 | 2 | 0.1 | 9 | 37.44 | 0.146611857 |
| 100110.1 | sp\|P22626\|ROA2_HUMAN | Heterogeneous nuclear ribonucleoproteins A2/B1 OS=Homo sapiens GN=HNRNPA2B1 PE=1 SV=2 | 2 | 0.1 | 37 | 21.39 | 0.146611857 |
| 100151.1 | sp\|P27348\|1433T_HUMAN | 14-3-3 protein theta OS=Homo sapiens GN=YWHAQ PE=1 SV=1 | 2 | 0.1 | 28 | 19.82 | 0.146611857 |
| 100169.1 | sp\|Q9BQE3\|TBA1C_HUMAN | Tubulin alpha-1C chain OS=Homo sapiens GN=TUBA1C PE=1 SV=1 | 2 | 0.1 | 50 | 8.02 | 0.146611857 |
| 100259.1 | sp\|P61981\|1433G_HUMAN | 14-3-3 protein gamma OS=Homo sapiens GN=YWHAG PE=1 SV=2 | 2 | 0.1 | 28 | 8.84 | 0.146611857 |
| 100646.1 | sp\|P67775\|PP2AA_HUMAN | Serine/threonine-protein phosphatase 2A catalytic subunit alpha isoform OS=Homo sapiens GN=PPP2CA PE=1 SV=1 | 2 | 0.1 | 36 | 2.81 | 0.146611857 |
| 100855.1 | sp\|P50990\|TCPQ_HUMAN | T-complex protein 1 subunit theta OS=Homo sapiens GN=CCT8 PE=1 SV=4 | 2 | 0.1 | 60 | 14.27 | 0.146611857 |
| 101227.1 | sp\|P21333\|FLNA_HUMAN | Filamin-A OS=Homo sapiens GN=FLNA PE=1 SV=4 | 2 | 0.1 | 281 | 12.65 | 0.146611857 |
| 101960.1 | sp\|Q9Y490\|TLN1_HUMAN | Talin-1 OS=Homo sapiens GN=TLN1 PE=1 SV=3 | 2 | 0.1 | 270 | 4.45 | 0.146611857 |
| 102769.1 | sp\|P55060\|XPO2_HUMAN | Exportin-2 OS=Homo sapiens GN=CSE1L PE=1 SV=3 | 2 | 0.1 | 110 | 6.34 | 0.146611857 |
| 105359.1 | sp\|P04271\|S100B_HUMAN | Protein S100-B OS=Homo sapiens GN=S100B PE=1 SV=2 | 2 | 0.1 | 11 | 9.34 | 0.146611857 |
| 106470.1 | sp\|Q9Y5L0\|TNPO3_HUMAN | Transportin-3 OS=Homo sapiens GN=TNPO3 PE=1 SV=3 | 2 | 0.1 | 104 | 2.88 | 0.146611857 |
| 100492.1 | sp\|P62826\|RAN_HUMAN | GTP-binding nuclear protein Ran OS=Homo sapiens GN=RAN PE=1 SV=3 | 6 | 12 | 24 | 118.81 | 0.153331729 |
| 111060.1 | sp\|Q8WZ42\|TITIN_HUMAN | Titin OS=Homo sapiens GN=TTN PE=1 SV=4 | 4 | 9 | 3814 | 0.22 | 0.16007254 |
| 100797.1 | sp\|O00410\|IPO5_HUMAN | Importin-5 OS=Homo sapiens GN=IPO5 PE=1 SV=4 | 41 | 30 | 124 | 116.96 | 0.190841486 |
| 102887.1 | sp\|P02786\|TFR1_HUMAN | Transferrin receptor protein 1 OS=Homo sapiens GN=TFRC PE=1 SV=2 | 16 | 10 | 85 | 63.08 | 0.237197244 |
| 103371.1 | sp\|P04844\|RPN2_HUMAN | Dolichyl-diphosphooligosaccharide--protein glycosyltransferase subunit 2 OS=Homo sapiens GN=RPN2 PE=1 SV=3 | 2 | 5 | 69 | 15.16 | 0.249110318 |
| 100945.1 | sp\|P68104\|EF1A1_HUMAN | Elongation factor 1-alpha 1 OS=Homo sapiens GN=EEF1A1 PE=1 SV=1 | 8 | 13 | 50 | 86.81 | 0.27292487 |
| 100167.1 | sp\|Q13509\|TBB3_HUMAN | Tubulin beta-3 chain OS=Homo sapiens GN=TUBB3 PE=1 SV=2 | 13 | 19 | 50 | 134.92 | 0.287415938 |
| 106288.1 | sp\|P27708\|PYR1_HUMAN | CAD protein OS=Homo sapiens GN=CAD PE=1 SV=3 | 6 | 3 | 243 | 5.56 | 0.312662745 |
| 101655.1 | sp\|P31943\|HNRH1_HUMAN | Heterogeneous nuclear ribonucleoprotein H OS=Homo sapiens GN=HNRNPH1 PE=1 SV=4 | 7 | 4 | 49 | 43.7 | 0.362666387 |
| 115909.1 | sp\|P55083\|MFAP4_HUMAN | Microfibril-associated glycoprotein 4 OS=Homo sapiens GN=MFAP4 PE=1 SV=2 | 19 | 14 | 29 | 242.74 | 0.383171688 |
| 100065.1 | tr\|E7EMB3\|E7EMB3_HUMAN | Calmodulin OS=Homo sapiens GN=CALM2 PE=1 SV=1 | 13 | 9 | 22 | 235.28 | 0.392451231 |
| 100959.1 | sp\|Q86VP6\|CAND1_HUMAN | Cullin-associated NEDD8-dissociated protein 1 OS=Homo sapiens GN=CAND1 PE=1 SV=2 | 5 | 8 | 136 | 30.08 | 0.403258826 |
| 100481.1 | sp\|P08133\|ANXA6_HUMAN | Annexin A6 OS=Homo sapiens GN=ANXA6 PE=1 SV=3 | 8 | 5 | 76 | 29.67 | 0.403258826 |
| 103127.1 | sp\|Q8N1F7\|NUP93_HUMAN | Nuclear pore complex protein Nup93 OS=Homo sapiens GN=NUP93 PE=1 SV=2 | 4 | 2 | 93 | 9.63 | 0.409725824 |
| 102085.1 | sp\|Q9BUF5\|TBB6_HUMAN | Tubulin beta-6 chain OS=Homo sapiens GN=TUBB6 PE=1 SV=1 | 6 | 9 | 50 | 84.29 | 0.43703108 |
| 103548.1 | sp\|Q14974\|IMB1_HUMAN | Importin subunit beta-1 OS=Homo sapiens GN=KPNB1 PE=1 SV=2 | 13 | 17 | 97 | 61.27 | 0.464543646 |
| 100202.1 | tr\|E9PJH4\|E9PJH4_HUMAN | 40S ribosomal protein S3 OS=Homo sapiens GN=RPS3 PE=1 SV=1 | 3 | 5 | 13 | 131.44 | 0.477161781 |
| 110235.1 | sp\|O14654\|IRS4_HUMAN | Insulin receptor substrate 4 OS=Homo sapiens GN=IRS4 PE=1 SV=1 | 5 | 3 | 134 | 14.21 | 0.477161781 |
| 100025.1 | tr\|Q96HY3\|Q96HY3_HUMAN | CALM1 protein OS=Homo sapiens GN=CALM3 PE=1 SV=1 | 11 | 8 | 13 | 345.79 | 0.490389144 |
| 100838.1 | sp\|Q00610\|CLH1_HUMAN | Clathrin heavy chain 1 OS=Homo sapiens GN=CLTC PE=1 SV=5 | 4 | 6 | 191 | 6.01 | 0.525692869 |
| 101587.1 | sp\|P01859\|IGHG2_HUMAN | Ig gamma-2 chain C region OS=Homo sapiens GN=IGHG2 PE=1 SV=2 | 5 | 7 | 36 | 104.52 | 0.562791465 |
| 107647.1 | sp\|Q12805\|FBLN3_HUMAN | EGF-containing fibulin-like extracellular matrix protein 1 OS=Homo sapiens GN=EFEMP1 PE=1 SV=2 | 6 | 8 | 55 | 39.37 | 0.592346844 |
| 100011.1 | sp\|P38646\|GRP75_HUMAN | Stress-70 protein, mitochondrial OS=Homo sapiens GN=HSPA9 PE=1 SV=2 | 44 | 49 | 74 | 141.24 | 0.604039106 |
| 102780.1 | sp\|P01860\|IGHG3_HUMAN | Ig gamma-3 chain C region OS=Homo sapiens GN=IGHG3 PE=1 SV=2 | 7 | 9 | 41 | 105.43 | 0.616614147 |
| 102211.1 | sp\|Q9Y265\|RUVB1_HUMAN | RuvB-like 1 OS=Homo sapiens GN=RUVBL1 PE=1 SV=1 | 2 | 3 | 50 | 25.9 | 0.653629236 |
| 102250.1 | sp\|Q00839\|HNRPU_HUMAN | Heterogeneous nuclear ribonucleoprotein U OS=Homo sapiens GN=HNRNPU PE=1 SV=6 | 11 | 9 | 91 | 37.56 | 0.654450842 |
| 108164.1 | sp\|Q9UBX5\|FBLN5_HUMAN | Fibulin-5 OS=Homo sapiens GN=FBLN5 PE=1 SV=1 | 4 | 3 | 50 | 22.93 | 0.704975995 |
| 101885.1 | sp\|P04632\|CPNS1_HUMAN | Calpain small subunit 1 OS=Homo sapiens GN=CAPNS1 PE=1 SV=1 | 15 | 13 | 28 | 189.05 | 0.705337391 |
| 100769.1 | sp\|O75340\|PDCD6_HUMAN | Programmed cell death protein 6 OS=Homo sapiens GN=PDCD6 PE=1 SV=1 | 5 | 4 | 22 | 59.48 | 0.738622707 |
| 102659.1 | sp\|P31689\|DNJA1_HUMAN | DnaJ homolog subfamily A member 1 OS=Homo sapiens GN=DNAJA1 PE=1 SV=2 | 7 | 6 | 45 | 57.98 | 0.781406072 |
| 103158.1 | sp\|P78527\|PRKDC_HUMAN | DNA-dependent protein kinase catalytic subunit OS=Homo sapiens GN=PRKDC PE=1 SV=3 | 12 | 11 | 469 | 21.86 | 0.834801667 |
| 101685.1 | sp\|P01834\|IGKC_HUMAN | Ig kappa chain C region OS=Homo sapiens GN=IGKC PE=1 SV=1 | 16 | 15 | 12 | 646.4 | 0.85745021 |
| 114762.1 | tr\|A0A087WZW8\|A0A087WZW8_HUMAN | Protein IGKV3-11 OS=Homo sapiens GN=IGKV3-11 PE=4 SV=1 | 16 | 15 | 26 | 298.99 | 0.85745021 |
| 100217.1 | tr\|K7ESA0\|K7ESA0_HUMAN | ATP synthase subunit alpha, mitochondrial OS=Homo sapiens GN=ATP5A1 PE=1 SV=1 | 2 | 2 | 8 | 30.26 | 1 |
| 101028.1 | sp\|P62979\|RS27A_HUMAN | Ubiquitin-40S ribosomal protein S27a OS=Homo sapiens GN=RPS27A PE=1 SV=2 | 3 | 3 | 18 | 69.62 | 1 |
| 101028.2 | tr\|J3QTR3\|J3QTR3_HUMAN | Ubiquitin (Fragment) OS=Homo sapiens GN=RPS27A PE=1 SV=1 | 3 | 3 | 12 | 155.34 | 1 |
| 101564.1 | sp\|Q99829\|CPNE1_HUMAN | Copine-1 OS=Homo sapiens GN=CPNE1 PE=1 SV=1 | 3 | 3 | 59 | 16.94 | 1 |
| 102279.1 | sp\|P17812\|PYRG1_HUMAN | CTP synthase 1 OS=Homo sapiens GN=CTPS1 PE=1 SV=2 | 4 | 4 | 67 | 26.26 | 1 |
| 100115.1 | sp\|P08758\|ANXA5_HUMAN | Annexin A5 OS=Homo sapiens GN=ANXA5 PE=1 SV=2 | 5 | 5 | 36 | 54.29 | 1 |
| 100481.3 | tr\|E5RK69\|E5RK69_HUMAN | Annexin OS=Homo sapiens GN=ANXA6 PE=1 SV=1 | 6 | 6 | 52 | 37.68 | 1 |
| 100831.1 | sp\|P28482\|MK01_HUMAN | Mitogen-activated protein kinase 1 OS=Homo sapiens GN=MAPK1 PE=1 SV=3 | 6 | 6 | 41 | 41.1 | 1 |
| 101913.1 | sp\|P01861\|IGHG4_HUMAN | Ig gamma-4 chain C region OS=Homo sapiens GN=IGHG4 PE=1 SV=1 | 6 | 6 | 36 | 83.52 | 1 |

**Table S2 Ubiquitination sites identified on KAT2A by mass spectrometry**

| Sequence | Modifications | Qvality PEP | Area: F686: Sample | Ions Score Mascot | Percolator PEP Mascot |
| --- | --- | --- | --- | --- | --- |
| QIPVESVPGIR |  | 0.0482975 | 4221323294 | 44 | 0.01638 |
| RGIIEFHVIGNSLTPK |  | 0.00122997 | 677278074.6 | 91 | 0.000142 |
| SEAPDYYEVIR |  | 0.0141228 | 8319565063 | 56 | 0.01151 |
| LFVADLQR |  | 0.0487187 | 1263354184 | 51 | 0.01662 |
| TLPENLTLEDAKR |  | 0.000535773 | 7113206173 | 73 | 0.000119 |
| TLPENLTLEDAKR | 1xGG [K12] | 0.00104814 | 347015782.3 | 71 | 0.000745 |
| ELKDPDQLYTTLK |  | 0.000666127 | 1592816116 | 59 | 6.45E-05 |
| ELKDPDQLYTTLK | 1xGG [K3] | 0.0108292 | 15557908.75 | 34 | 0.002351 |
| RTLILTHFPK |  | 0.0133928 | 23397616 | 33 | 0.003102 |
| DPDQLYTTLK |  | 0.00956593 | 588248927 | 51 | 0.002006 |
| IPYTELSHIIK |  | 0.00205024 | 6508706533 | 58 | 0.000275 |
| SHPSAWPFMEPVK | 1xOxidation [M9] | 0.00713915 | 514624347.9 | 39 | 0.01257 |
| IPYTELSHIIKK |  | 0.00129733 | 664386396.7 | 61 | 0.000152 |
| TLPENLTLEDAK |  | 0.00673911 | 5302138839 | 50 | 0.001274 |
| SEAPDYYEVIRFPIDLK |  | 0.00237397 | 317811366.1 | 55 | 0.002043 |
| LLGMVVDVENLFMSVHK | 2xOxidation [M4; M13] | 0.000663174 | 965343419 | 60 | 6.42E-05 |
| KSEAPDYYEVIRFPIDLK |  | 0.00641814 | 20811887.81 | 29 | 0.001197 |
| KCILQMTRPVVEGSLGSPPFEKPNIEQGVLNFVQYK | 1xCarbamidomethyl [C2]; 1xOxidation [M6] | 0.0016273 | 125789086 | 54 | 0.000204 |
| VMGDIPMELVNEVMLTITDPAAMLGPETSLLSANAAR |  | 0.00123545 | 5896319.5 | 38 | 0.000143 |
| GIIEFHVIGNSLTPK |  | 0.00156351 | 1910503558 | 73 | 0.000193 |
| QVYFYLFK |  | 0.0489307 | 2078427956 | 41 | 0.01668 |
| LLGMVVDVENLFMSVHKEEDTDTK | 2xOxidation [M4; M13] | 4.36827E-05 | 3313442486 | 105 | 2.15E-06 |
| LLGMVVDVENLFMSVHKEEDTDTK | 1xGG [K17]; 2xOxidation [M4; M13] | 0.00440125 | 7563530 | 33 | 0.000735 |
| EEDTDTKQVYFYLFK |  | 0.00895069 | 118045113.8 | 39 | 0.00345 |
| KSEAPDYYEVIR | 1xGG [K1] | 0.00357226 | 52085800.5 | 67 | 0.000563 |
| KSEAPDYYEVIR |  | 0.000387326 | 180893357.5 | 70 | 3.21E-05 |
| ERQTMFELSK | 1xOxidation [M5] | 0.0623364 | 23149561.06 | 41 | 0.08643 |
| YETTHVFGR |  | 0.00973699 | 2446437751 | 66 | 0.00205 |
| ETGWKPLGK |  | 0.0386787 | 2258591804 | 36 | 0.02684 |
| SQAEDVATYKVNYTR |  | 0.000189368 | 201525244.4 | 87 | 1.27E-05 |
| SQAEDVATYKVNYTR | 1xGG [K10] | 0.00299083 | 10683778.75 | 46 | 0.000446 |
| QTMFELSK | 1xOxidation [M3] | 0.0186554 | 776275455 | 52 | 0.004747 |
| TLALIKDGR |  | 0.0653569 | 123081194.6 | 51 | 0.02442 |
| SQAEDVATYK |  | 0.0163407 | 3267001081 | 49 | 0.004007 |
| GYGTHLMNHLK | 1xOxidation [M7] | 0.0206474 | 551361523.3 | 32 | 0.005438 |
| MPKEYIAR | 1xOxidation [M1] | 0.0438916 | 17424596.88 | 39 | 0.01447 |
| NPKPPTAPR | 1xGG [K3] | 0.0189876 | 3780258.984 | 48 | 0.00486 |
| NPKPPTAPR |  | 0.00704478 | 150398361.5 | 51 | 0.001348 |
| HKTLALIKDGR |  | 0.00895069 | 13605827.75 | 68 | 0.001837 |
| RTLPENLTLEDAKR |  | 0.00458066 | 42800277.5 | 60 | 0.000777 |
| LETPAQFR |  | 0.0667759 | 7726837899 | 51 | 0.06701 |
| KLFVADLQR |  | 0.0224506 | 468781694 | 46 | 0.01077 |
| SHPSAWPFMEPVKK | 1xOxidation [M9] | 0.00103886 | 2539015890 | 50 | 0.000114 |
| SHPSAWPFMEPVKK | 1xGG [K]; 1xOxidation [M9] | 0.0213885 | 30720736.25 | 24 | 0.0185 |
| GKELKDPDQLYTTLK |  | 0.000331526 | 380412838.5 | 63 | 0.000161 |
| GKELKDPDQLYTTLK | 1xAcetyl [N-Term] | 0.00118175 | 1254802903 | 58 | 0.000562 |
| GKELKDPDQLYTTLK | 1xGG [K2] | 0.0118845 | 1946268.609 | 39 | 0.002655 |
| TLILTHFPK |  | 0.0736922 | 6727419746 | 41 | 0.03144 |
| ETGWKPLGKEK | 1xGG [K9] | 0.0775673 | 36355214.5 | 22 | 0.03044 |
| LKEGGLIDK |  | 0.0778988 | 635666569.8 | 38 | 0.03065 |
| LKEGGLIDK | 1xGG [K2] | 0.107859 | 5426602.375 | 16 | 0.04718 |
| GKELKDPDQLYTTLK | 1xAcetyl [N-Term]; 1xGG [K5] | 0.118707 | 13336458 | 22 | 0.05349 |
| KSEAPDYYEVIR | 1xAcetyl [N-Term] | 0.132169 | 3656305.563 | 34 | 0.06171 |
| SEAPDYYEVIRFPIDLKTMTER | 1xOxidation [M19] | 0.136582 | 32074800.03 | 7 | 0.0645 |
| FSHLAPR |  | 0.134911 | 2817443512 | 36 | 0.06363 |
| TLPENLTLEDAKRLR | 1xGG [K12] | 0.162648 | 33403984 | 18 | 0.08174 |
| NLLAQIK |  | 0.178319 | 2541056032 | 37 | 0.1007 |
| DKLVPEKR |  | 0.184799 | 315329248 | 39 | 0.09707 |
| SRYYVTR |  | 0.179742 | 8606736.313 | 29 | 0.09354 |
